# Evidence of positive and negative selection associated with DNA methylation

**DOI:** 10.1101/2021.11.25.469994

**Authors:** Charlie Hatcher, Genetics of DNA Methylation Consortium, Gibran Hemani, Santiago Rodriguez, Tom R. Gaunt, Daniel J. Lawson, Josine L. Min

## Abstract

Signatures of negative selection are pervasive amongst complex traits and diseases. However, it is unclear whether such signatures exist for DNA methylation (DNAm) that has been proposed to have a functional role in disease. We estimate polygenicity, SNP-based heritability and model the joint distribution of effect size and minor allele frequency (MAF) to estimate a selection coefficient (*S*) for 2000 heritable DNAm sites in 1774 individuals from the Avon Longitudinal Study of Parents and Children. Additionally, we estimate *S* for meta stable epi alleles and DNAm sites associated with aging and mortality, birthweight and body mass index. Quantification of MAF-dependent genetic architectures estimated from genotype and DNAm reveal evidence of positive (*S* > 0) and negative selection (*S* < 0) and confirm previous evidence of negative selection for birthweight. Evidence of both negative and positive selection highlights the role of DNAm as an intermediary in multiple biological pathways with competing function.

## Introduction

Genome-wide association studies (GWASs) have identified many genetic variants (single nucleotide polymorphisms; SNPs) associated with complex traits and diseases^1^. Natural selection plays a role in influencing the genetic architecture of complex traits, altering allele frequency at many genetic loci^2^. Negative selection prevents deleterious mutations from becoming common^3^ and is thought to explain why GWASs have identified many common variants of low effect size^4^. Several studies have shown evidence of negative selection acting on complex traits (including height, body mass index; BMI and birthweight) using the relationship between minor allele frequency (MAF) and SNP effect size to estimate a selection coefficient (*S*)^3,5,6^. However, there is difficulty in separating the action of selection from genetic drift when using MAF and SNP effect size to characterise genetic architecture^7^.

Most GWA loci reside in non-coding regions and colocalization studies have shown that genetic factors underlying intermediate traits are shared with GWA loci^8,9^. Intermediate traits such DNA methylation (DNAm) and gene expression may therefore also show signatures of selection. Variation in DNAm can be influenced by age^10^, environmental^11^, genetic^12^ and stochastic^13^ changes. The variability of DNAm maybe caused by natural selection, epigenetic stochasticity^14^ or cellular plasticity^15^. The Genetics of DNA Methylation Consortium (GoDMC) has identified a large number of methylation quantitative trait loci (mQTLs) in blood^16^. They showed that these DNAm sites influenced by genetic factors are polygenic^16^. mQTLs were enriched for a variety of selection metrics (including the singleton density score; SDS^17^ and fixation index; *F_st_* ^18^) and show a strong negative relationship between MAF and mQTL effect size^19^. It is therefore likely that natural selection acts on many mQTL variants jointly. However, selection is difficult to detect as DNAm is typically controlled by a local *cis* variant with large effect size and many physically separated *trans* variants with small effect sizes. Previous studies on *cis*-regulatory regions have found evidence of purifying selection on sequence-dependent allele-specific DNAm^20^ and positive selection among African agriculturist populations^21^. Similarly, gene expression traits are polygenic^22^ and SNPs showing signatures of selection are enriched among SNPs associated with gene expression (expression quantitative trait loci; eQTLs)^3,23^.

DNAm has a variety of roles in gene regulation^24,25^, is likely cell type-specific, and can be used as a biomarker for risk stratification and disease detection^26,27^. DNAm at cytosine-guanine dinucleotides (CpGs) has been associated with repression of transcription factor (TF) binding, however, TF binding has also been shown to inhibit DNAm^28^. Across the 450k sites most commonly measured in epidemiological studies^29^ (which are biased to promoter regions), mean heritability for DNAm has been shown to be around 20%^12^ and relationships between the heritability of a DNAm site and the number of mQTLs and between heritability and effect size have been found^16^ DNAm sites may have particular properties in terms of natural selection where heritable sites should have increased polygenicity with a larger proportion of SNPs with larger effect sizes^4^. In epigenome-wide association studies (EWASs), DNAm sites have been associated with many complex traits and diseases including those showing signatures of negative selection such as BMI and birthweight^3,30,31^. Additionally, PhenoAge is a composite DNAm predictor of aging (trained on mortality including 42 clinical measures and age), that has been predictive of disease risk and mortality^32^. To date, there is little known about whether these sites are a target of selection for example due to antagonistic pleiotropy^33^ where genes required for earlier stages of development may have deleterious effects in later life^32^. Meta stable DNAm sites exhibiting greater similarity than can be explained genetically have also been identified^14^. It may be the case that increased variability of these sites occurred as initial response to the environment before the effect of natural selection.

DNAm may play various roles in underlying biological processes, and therefore we expect it to be subject to both positive and negative selection. Here, we investigate the relationship between MAF and effect size for SNPs at individual DNAm sites from the widely used 450k array to make inferences about the action of natural selection, which we hypothesise may vary for each DNAm site. We utilise *BayesNS,* a Bayesian mixed linear model method (MLM) that estimates polygenicity, SNP-based heritability and the joint distribution of MAF and effect size^3^. We apply *BayesNS* to DNAm data from the Accessible Resource for Integrated Epigenomic Studies (ARIES) cohort^34^.

## Results

### Estimation of genetic architecture parameters of DNAm sites

We used a Bayesian mixed linear model (*BayesNS*) to estimate genetic architecture parameters of DNAm sites including polygenicity, SNP-based heritability and a selection coefficient (*S*)^3^. We applied *BayesNS* to DNAm sites profiled in blood from 1774 mother-offspring individuals from ARIES^34^ and 474,939 independent non-major histocompatibility complex (MHC) and non-lactase (LCT) SNPs. Specifically, we considered 2000 DNAm sites which have ‘high’ heritability estimates from twin studies (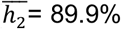, range 79-99%, Table 1)^12^, as selection is dependent on a genetic contribution to DNAm variance. Secondly, we analysed 1508 DNAm sites which show non-genetically mediated similarity between monozygotic twins, so-called epigenetic supersimilarity (ESS) DNAm sites^14^. Blood DNA methylation at ESS DNAm sites exhibit plasticity to the periconceptional environment and is associated with risk of cancer. Finally, we considered 513 DNAm sites which combined predict biological age (“PhenoAge”), a trait that is moderately heritable and has been associated with aging, mortality and is predictive of cardiovascular disease risk^32^.

**Table 1.**
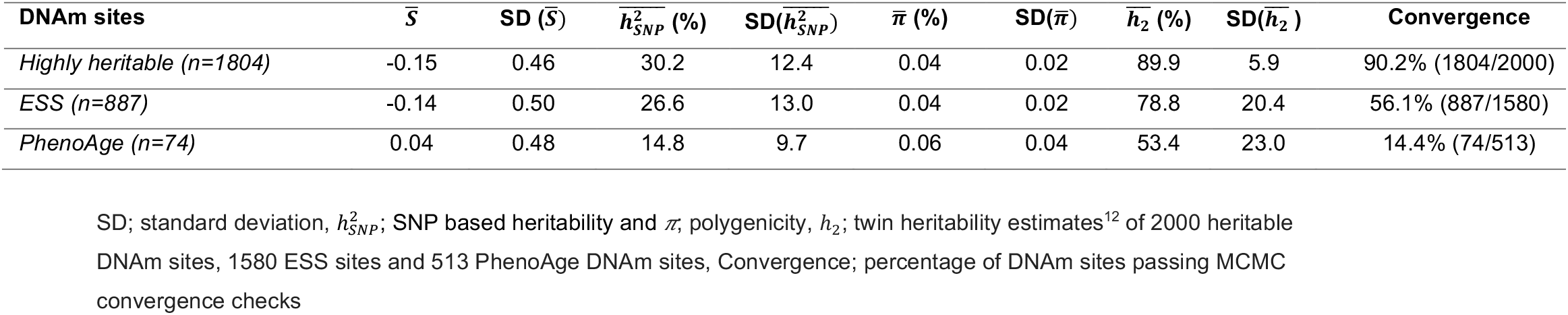
Estimation of genetic architecture parameters for highly heritable, ESS and PhenoAge DNAm sites.

Convergence of the Markov chain Monte Carlo (MCMC) algorithm implies that a single consistent selection signal is found, whilst failure to converge implies that competing, inconsistent sets of SNPs explain the data equally (and poorly). This was assessed with the Raftery-Lewis long-chain diagnostic test^35^ and MCMC trace plots (Figures S1;S2;S3). In line with previous work^3^, DNAm sites which failed convergence checks typically had lower estimates of heritability (Figure S3; Table 1). *BayesNS* estimates SNP-based heritability and as with the estimates of twin heritability, the highly heritable DNAm sites had the highest mean estimate of SNP based heritability (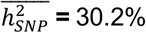; SD=12.4%; Table 1), followed by the ESS DNAm sites (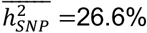; SD=13.0%; Table 1) and then the PhenoAge DNAm sites (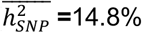; SD=9.7%;Table 1). Since we only consider DNAm sites which passed MCMC convergence diagnostics, (Table 1; Figure S1-S3), these mean estimates are likely higher than we would expect for each set of DNAm sites.

### DNAm shows signatures of both positive and negative selection

*BayesNS* uses the relationship between SNP effect size and MAF to estimate a selection coefficient (*S*)^3^. When *S* = 0 effect size is independent of MAF and this would reflect a ‘neutral’ scenario, an *S* > 0 would represent evidence of positive selection and an *S* < 0 would represent evidence of negative selection. Quantification of MAF-dependent genetic architectures revealed the action of both positive *S* > 0 and negative *S* < 0 selection across all three sets of DNAm sites (Figure 1). On average, estimates are close to zero, (PhenoAge DNAm sites; 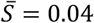) being mildly negative for the highly heritable 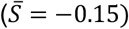 and ESS DNAm sites 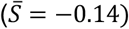 (Table 1). Across the distributions we see individual DNAm sites with more extreme positive and negative values of S. DNAm sites with extreme negative estimates of *S* (*S* < −1) are annotated to a variety of genes including those involved in transcription (*ATF7IP)^36^* and tumour suppression (*SCRIB*)^37^. DNAm sites with extreme positive estimates of *S* (*S* > 1) are annotated to a variety of genes including cg07175007 (*S*=1.13, SD=0.56) near *UHMK1* associated with the cell cycle^38^ and cg0479814 (*S*=0.65, SD=0.60) near *SMYD3* a histone methyltransferase ^39^(Table S1).

**Figure 1.**
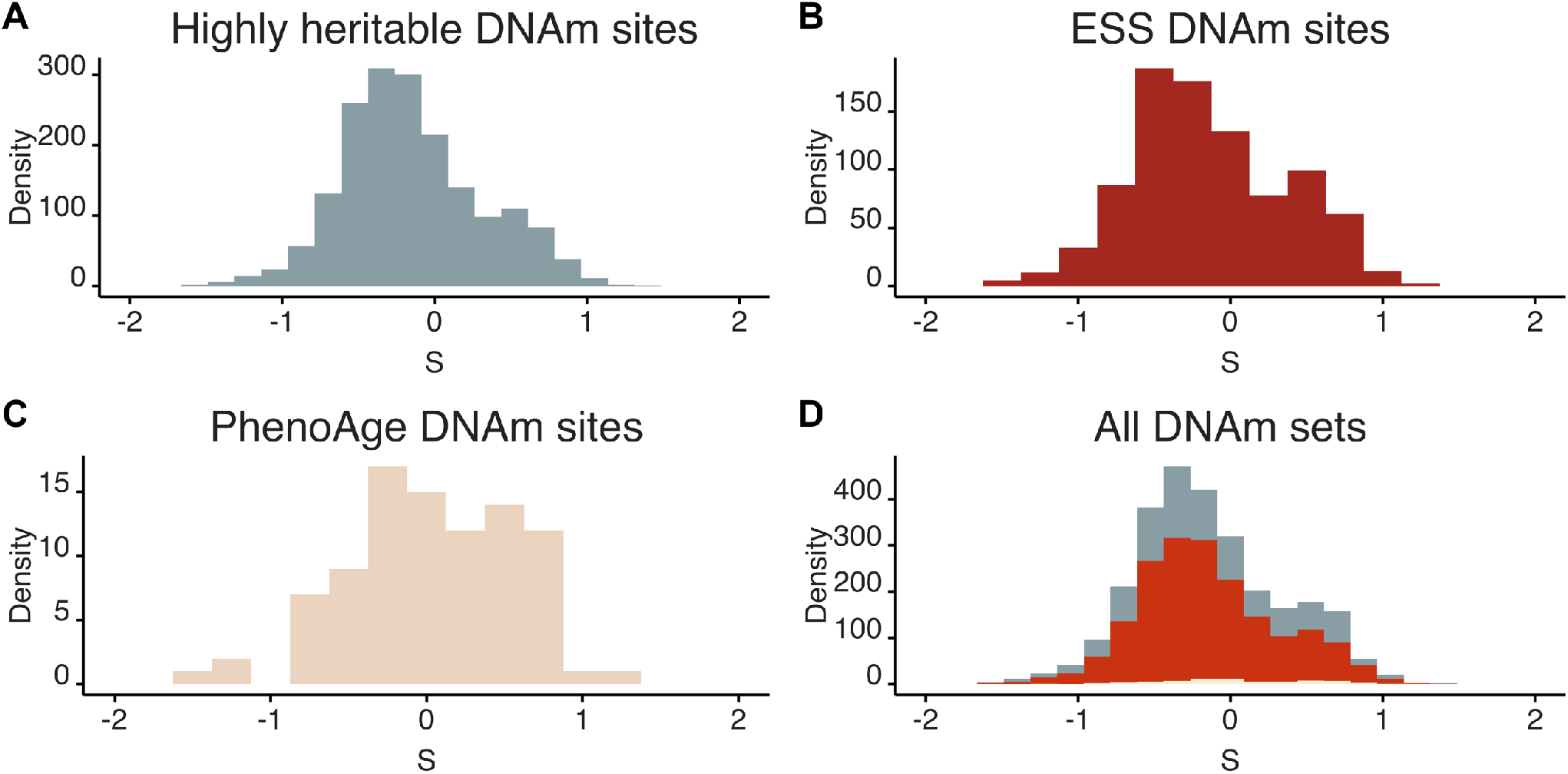
Estimates of *S* from *BayesNS*. *S* is estimated using the relationship between SNP effect size and MAF, when *S* = 0 SNP effect size is independent of MAF (neutral), *S* > 0 indicates positive selection, *S* < 0 indicates negative selection. Results are for **(A)** 1804 highly heritable DNAm sites, **(B)** 887 ESS DNAm sites and **(C)** 74 PhenoAge DNAm sites which passed MCMC convergence checks. All DNAm sites shown in **(D)** with highly heritable DNAm sites in grey, ESS DNAm sites in red and PhenoAge DNAm sites in beige.

As a sensitivity analysis we implemented models to account for genetic drift^7^ (Figure S4), which suggest that drift may be important but is not the sole driver of the signal of selection, supporting the hypothesis that this captures real biological processes. However, we cannot rule out that any specific effect was not caused by genetic drift.

### Polygenicity is associated with selection

*BayesNS* estimates polygenicity as the proportion of 200kb genomic ‘windows’ with non-zero effects^3^. In contrast to findings from GoDMC^19^, our results suggest that the genetic architecture of DNAm is not very polygenic (highly heritable DNAm sites: 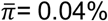, ESS DNAm sites 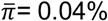, PhenoAge DNAm sites: 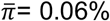; Table 1). This finding is in part due to the bimodality of the effect size distribution in DNAm: we lack power to capture polygenic *trans* mQTLs with low effect sizes, whilst we are powered to detect large *cis* mQTL effects. However, it may also reflect the role of many DNAm sites in biological pathways, having a specific biological purpose but either affecting, or being affected by, many other processes.

We additionally investigated the number of SNPs (N SNPs) highly associated with each DNAm site (posterior inclusion probability; *PIP* >=0.8). Across all three sets of DNAm sites, we find a negative relationship between S and N SNPs for DNAm sites (regression coefficient for highly heritable DNAm sites; −0.09; *p* < 2.2 ×10^−16^, ESS DNAm sites; −0.10; *p* < 2.2 × 10^−16^, PhenoAge DNAm sites; −0.14; *p* = 0.00064; Figure 2). In addition, we find that SNPs associated with DNAm sites with negative estimates of *S* have lower mean estimates of variance explained (VE) compared to those with positive estimates of *S* (Figure 3). Polygenicity is therefore associated with selection, with DNAm associated with few mQTLs being the only class of positive selection, and highly polygenic DNAm being subject to strictly negative selection. Further, positively selected DNAm tends to have almost all of the heritability accounted for by identifiable mQTLs.

**Figure 2.**
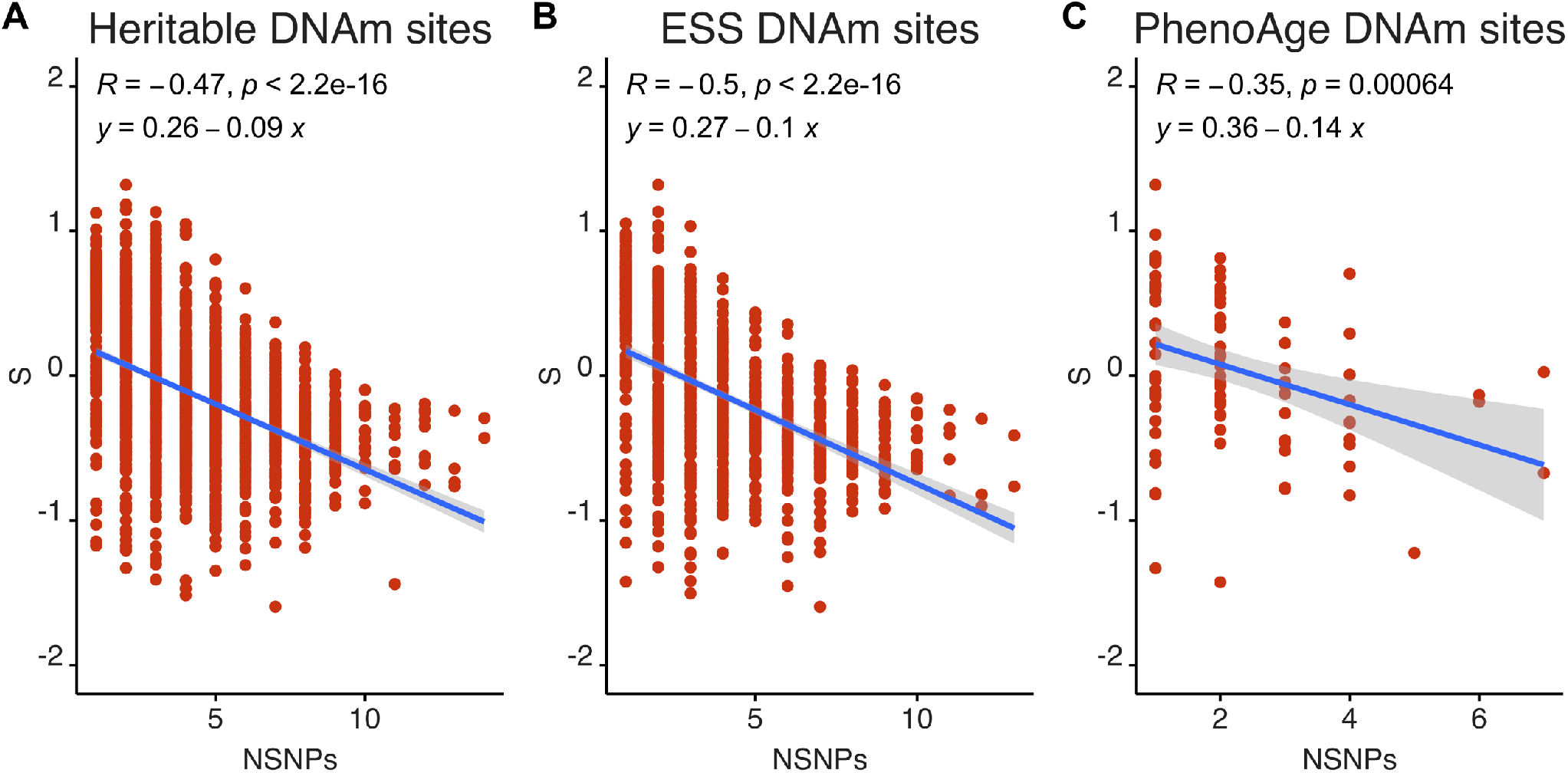
Relationship between estimates of *S* from *BayesNS* and NSNPs. For **(A)** 1804 highly heritable DNAm sites, **(B)** 887 ESS DNAm sites and **(C)** 74 PhenoAge DNAm sites. NSNPs calculated as the number of SNPs with a posterior inclusion probability (PIP) >=0.8 for each DNAm site and *S* calculated from the relationship between SNP effect size and MAF. Slope for highly heritable DNAm sites (−0.09; *p* < 2.2 ×10^−16^), ESS DNAm sites (−0.10; *p* < 2.2 ×10^−16^), PhenoAge DNAm sites (−0.14; *p* = 0.00064).

**Figure 3:**
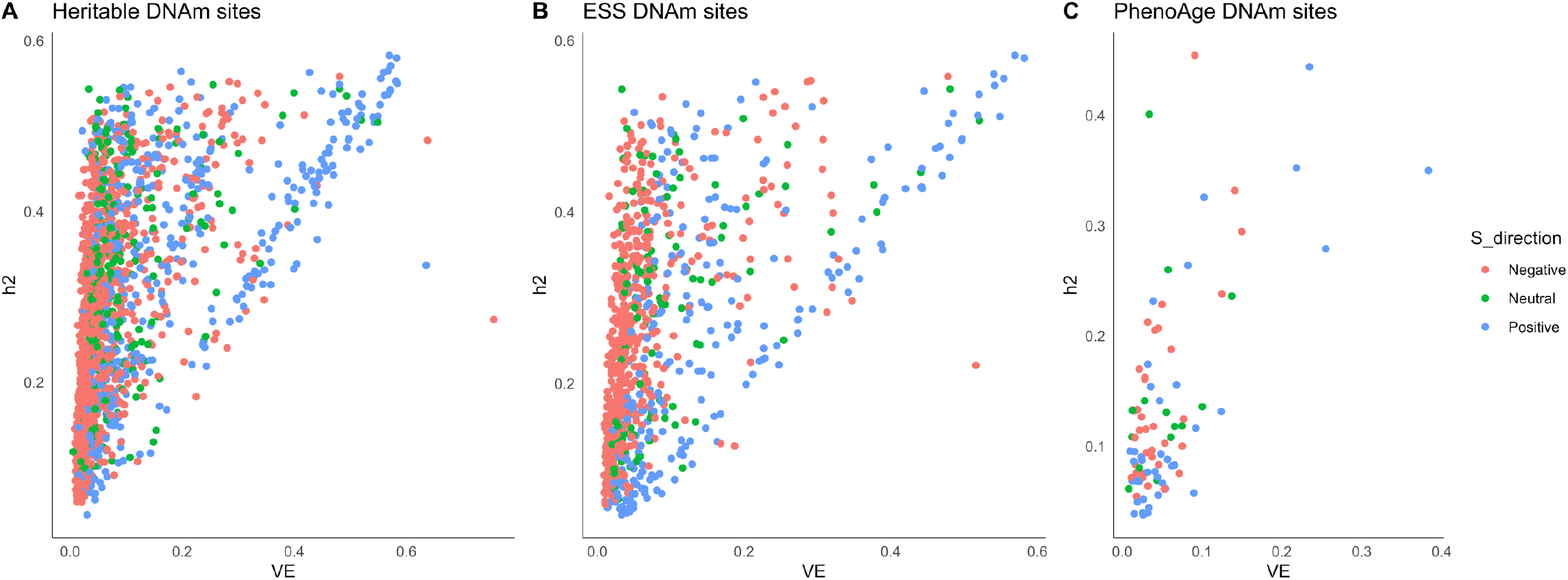
Relationship between h^2^_SNP_ calculated by *BayesNS* and mean variance explained (VE). For **(A)** 1804 highly heritable DNAm sites, **(B)** 887 ESS DNAm sites and **(C)** 74 PhenoAge DNAm sites. Estimates of *S* coloured: red (negative <=−0.1), green (neutral −0.1 – 0.1) and blue (positive >0.1).

### Relationship between selection estimates and traditional selection measures

We additionally investigated whether estimates of S correlate with five selection metrics: SDS^17^, *F_st_*^40^, integrated haplotype score (iHS)^41^, cross-population extended haplotype homozygosity XPEHH^42^ (CEU v. YRI) and XPEHH (CEU v. CHB) in sets of ‘high’ *PIP* (*PIP* > 0.1) and ‘all’ *PIP* (*PIP* >= 0.001) SNPs for each DNAm site. Values of *S* have the highest correlation with *F_st_*^40^(0.193; Figure S5A, 0.113; Figure S5C, 0.151; Figure S5E, for highly heritable, ESS and PhenoAge DNAm sites respectively), however, when we include ‘all’ possible SNPs, even though we weight by *PIP,* the correlation becomes negative and tends to decrease in magnitude (−0.045; Figure S5B, −0.127; Figure S5D, −0.091; Figure S5F). This implies that *cis* or strongly acting *trans* SNPs are selected differently to the bulk DNA associations, i.e. that they are selected via a different mechanism, and that the low *PIP* SNPs are subject to a diversity of pathways, hence leading to an average selection close to 0 (Table 1). We additionally calculate correlations between S and LD scores ^43^. Correlations between LD scores and *S* are small, with the lowest magnitude correlation being −0.005 and the highest being −0.065, suggesting that estimates of *S* are not correlated with LDSC.

### Biological properties of DNAm sites under selection

The magnitude of *S* is related to the ‘strength of selection on trait-associated SNPs’^3^. To understand whether DNAm traits under ‘stronger’ selection had biological relevance, we assessed whether DNAm sites with estimates of S ≤ −0.5 and ≥ 0.5 were enriched or depleted for predicted chromatin states^44^. The positive highly heritable DNAm sites (n=212, *S* ≥ 0.5;) showed the strongest enrichment (qvalue<0.05) for enhancers (Odds Ratio; ORs EnhW1 1.75-3.21; ORs EnhW2 1.72-2.57) and promoters (PromP ORs=1.84-3.26; PromU ORs=1.52-2.18) (Figure 4A; Table S2). The negative highly heritable DNAm sites (n=376, *S* ≤ −0.5; Figure 4B; Table S2) showed only enrichment for poised promoters (PromP, OR=2.1) but not for transcription activity. Both positive (n=123, *S* ≥ 0.5) and negative (n=218, *S* ≤ −0.5) ESS DNAm sites also show enrichment for poised promoters (PromP) across all tissue types (positive ORs 2.24-5.1 negative ORs: 1.79-3.64; Figure S7; Tables S3-S4). Poised chromatin is associated with both activating and repressing histone modifications and has been proposed to play a role in the prevention of DNAm^45^. DNAm sites showing signatures of selection are therefore enriched for bivalent chromatin structure associated with silencing genes whilst keeping them ready for activation^46^. CpG rich promoters have been shown to be subject to ‘epigenetic buffering’ against the effects of random mutations due to their association with housekeeping genes^20^.

**Figure 4:**
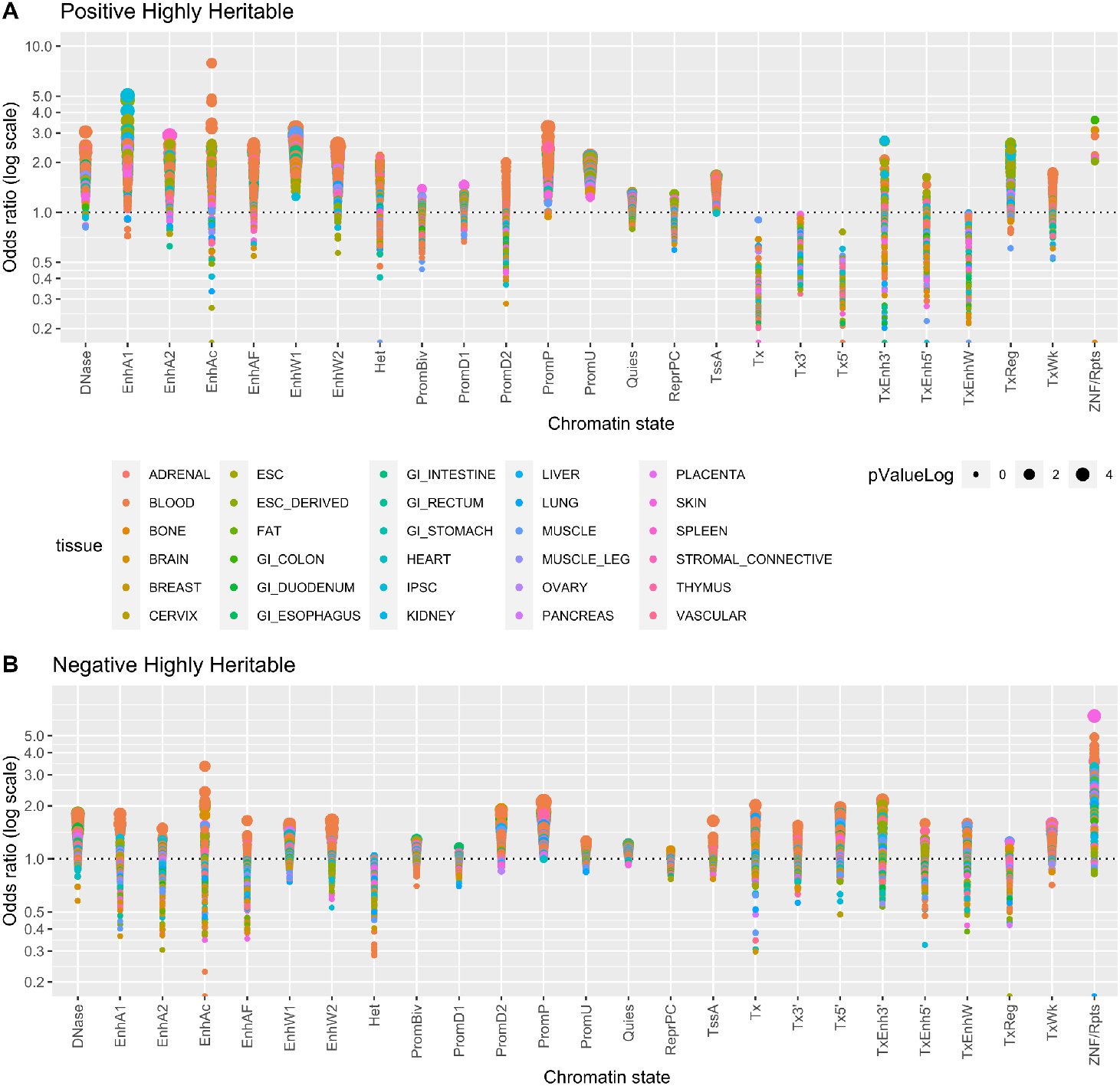
Enrichment or depletion of DNAm sites in predicted chromatin states for DNAm sites from the highly heritable probe set with estimates of *S* >=0.5 (positive highly heritable) and *S* <= −0.5 (negative highly heritable). Odds ratio (on log scale) shown on the *y* axis and chromatin state on the *x* axis. Size of circle represents the −log10 *P* value. Enrichment analysis performed via two-sided Fisher’s exact test implemented in LOLA^1^. 25 chromatin states abbreviations: TssA, Active TSS; PromU, Promoter Upstream TSS; PromD1, Promoter Downstream TSS with DNase; PromD2, Promoter Downstream TSS; Tx5’, Transcription 5’; Tx, Transcription; Tx3’, Transcription 3’; TxWk, Weak transcription; TxReg, Transcription Regulatory; TxEnh5’, Transcription 5’ Enhancer; TxEnh3’, Transcription 3’ Enhancer; TxEnhW, Transcription Weak Enhancer; EnhA1, Active Enhancer 1; EnhA2, Active Enhancer 2; EnhAF, Active Enhancer Flank; EnhW1, Weak Enhancer 1; EnhW2, Weak Enhancer 2; EnhAc, Enhancer Acetylation Only; DNase, DNase only; ZNF/Rpts, ZNF genes & repeats; Het, Heterochromatin; PromP, Poised Promoter; PromBiv, Bivalent Promoter; ReprPC, Repressed PolyComb, Quies, Quiescent/Low.

**Figure 5:**
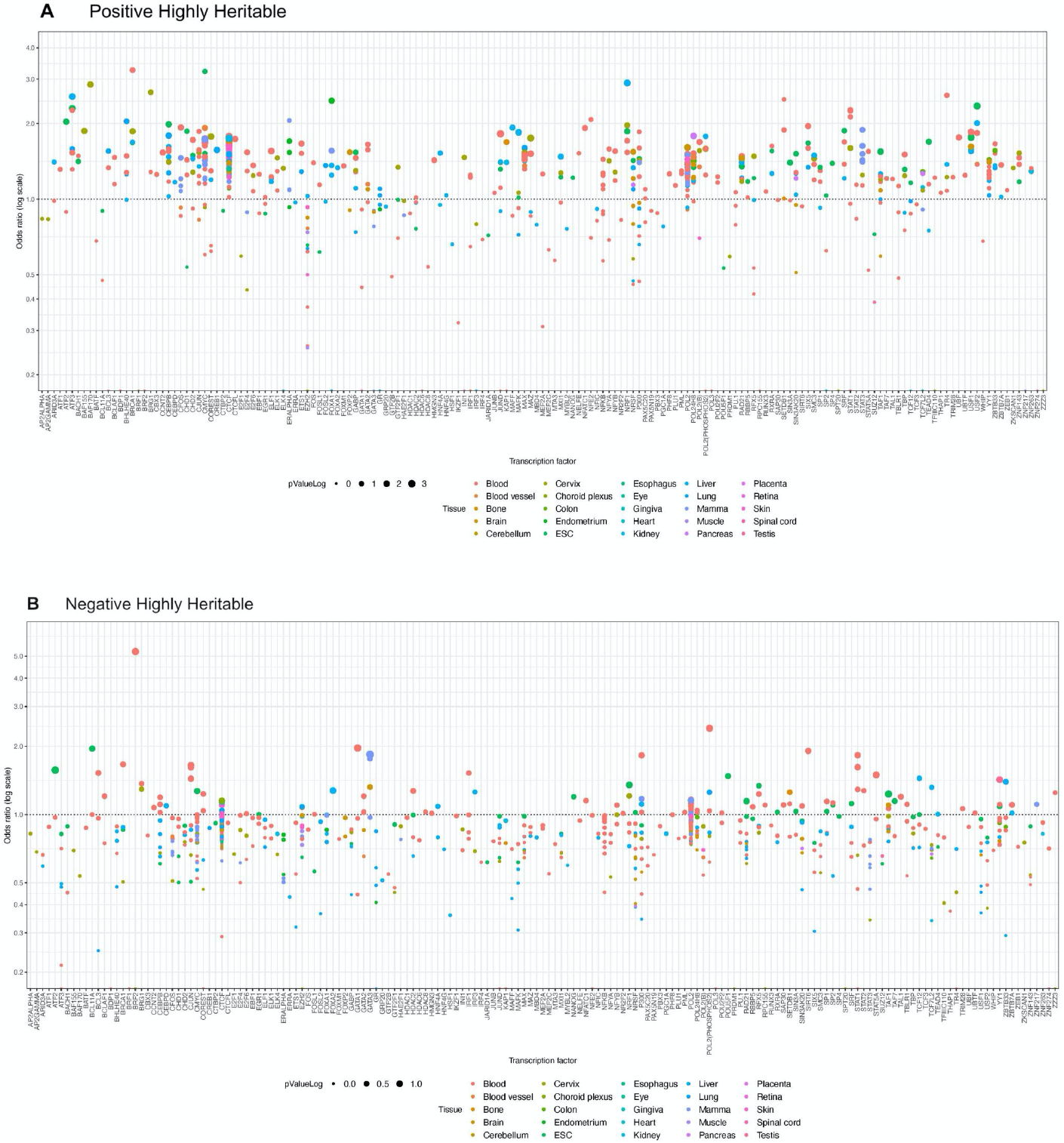
Enrichment or depletion of DNAm sites in transcription factors (TFs) for DNAm sites from the highly heritable probe set with estimates of *S* > = 0.5 (positive highly heritable) and *S* <= −0.5 (negative highly heritable). Odds ratio (on log scale) shown on the *y* axis and chromatin state on the *x* axis. Size of circle represents the –log10 *P* value. Enrichment analysis performed via two-sided Fisher’s exact test implemented in LOLA^1^.

We additionally assessed enrichment of 167 transcription factor binding sites (TFBSs) in 127 different cell types comprising 30 tissues^47^. Transcription factors have previously been shown to be under weak purifying selection, with a limited minority exhibiting signatures of positive selection^48^. Arbiza *et al.,* find evidence of positive selection on GATA-binding zinc finger proteins^48^. Though, we do not see evidence of enrichment for TFBS for our DNAm sites of interest (Tables S6-S9).

### BMI and birthweight associated DNAm sites show signatures of selection

We used *BayesNS* to estimate *S*, 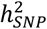 and *π* for 893 DNAm sites associated with birthweight^31^ and 243 DNAm sites associated with BMI^30^ in EWAS of individuals of European ancestry^49^. 220 birthweight-associated DNAm sites (24.6%) and 42 BMI-associated DNAm sites (17.0%) passed MCMC convergence tests (Figure S3). As with the highly heritable, ESS and PhenoAge DNAm sites, MAF-dependent genetic architectures estimated from genotype and DNAm revealed the action of both positive and negative selection for BMI (Š; 0.04; SD; 0.59, range: −1.09:0.98) and birthweight (Š; −0.05, SD; 0.59, range: −1.81:0.98) associated DNAm sites (Figure 6; Table S10). Birthweight-associated cg16875057 has an *S* estimate of −1.81 and is annotated to the *STK39* gene which is associated with the cellular stress response pathway and hypertension^50^. In addition, birthweight-associated cg07157107 (S=0.98) is associated with the nicotinic receptor *CHRNA6,* positive selection has previously been reported on genomic regions containing nicotinic receptor genes^51^. In contrast, a previous study using BMI and birthweight GWA loci found only evidence of negative selection^3^. After adjustment for non-random properties of the DNAm sites, we found that birthweight associated DNAm sites showed an enrichment of negative estimates of *S* as compared to heritability matched background DNAm sites (Table 2). To assess whether biological pathways were enriched among the DNAm sites with extreme *S* we performed GOterm enrichment analysis, however none of the pathways showed evidence of enrichment.

**Figure 6:**
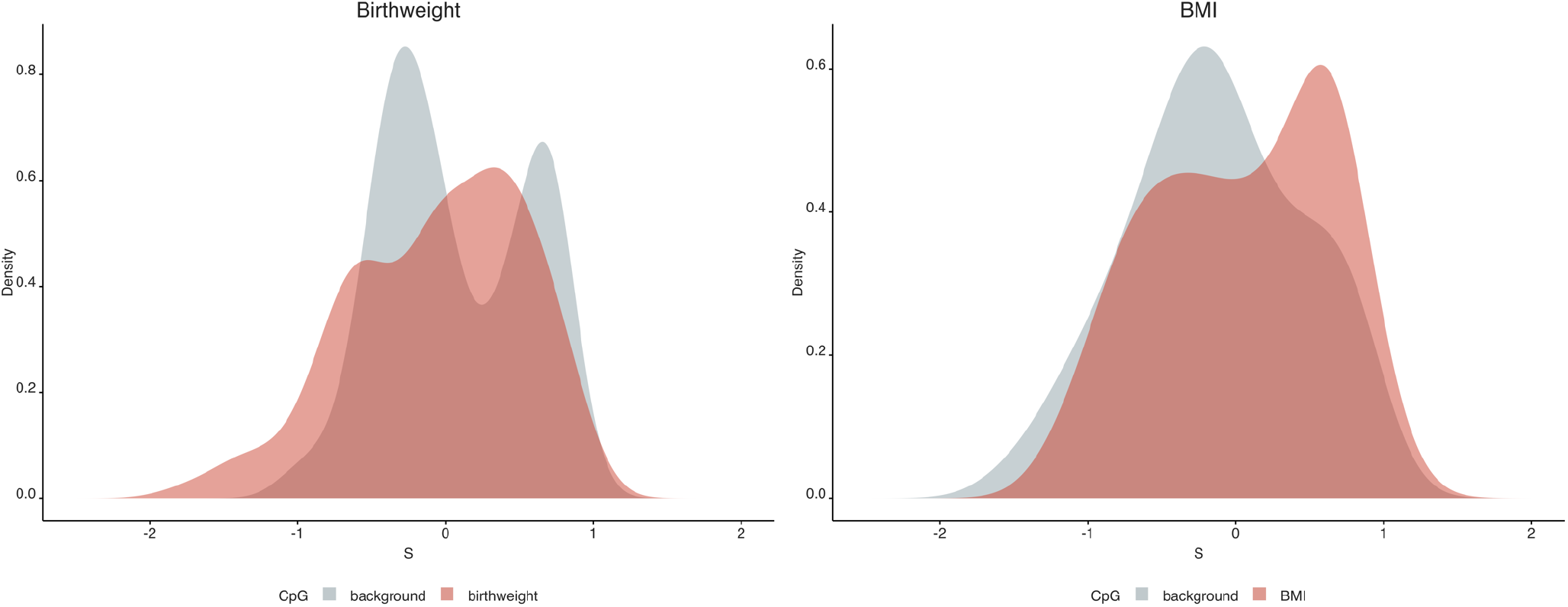
Distribution of estimates of *S* for DNAm sites associated with birthweight and BMI compared to background DNAm sites. *S* is estimated using the relationship between SNP effect size and MAF, when *S* = 0 SNP effect size is independent of MAF (neutral), *S* > 0 indicates positive selection, *S* < 0 indicates negative selection. Birthweight and BMI associated DNAm sites shown in red and matched DNAm sites (matched on GC/CpG content and heritability) shown in grey.

**Table 2.** Birthweight-associated DNAm sites are enriched for negative estimates of *S*.

| DNAm sites | Fisher's exact test $P$-value | Odds ratio (OR) | Lower 95% CI | Upper 95% CI |
| --- | --- | --- | --- | --- |
| *PhenoAge* ($n=74$) | 1 | 1.09 | 0.44 | 2.70 |
| *BMI-associated* ($n=42$) | 1 | 1.13 | 0.37 | 3.45 |
| *Birthweight* ($n=220$) | $3.432 \times 10^{-6}$ | 0.29 | 0.16 | 0.50 |

## Discussion

In this study, we have characterized the genetic architecture of DNA methylation at individual DNAm sites measured on the 450k array^29^. Specifically, we consider estimates of polygenicity, SNP-based heritability and the joint-distribution of effect size and MAF for 1804 highly heritable DNAm sites, 887 ESS DNAm sites and 74 PhenoAge DNAm sites. Unlike previous work looking at complex traits and gene expression which find evidence of negative selection exclusively^3,5,6^, across all sets of DNAm sites we find evidence of both positive (*S* > 0) and negative selection (*S* < 0). These findings support previous research showing an enrichment of mQTLs among SNPs with signatures of positive selection, plus a negative relationship between MAF and mQTL effect size^19^. We were able to estimate *S* at individual DNAm sites allowing us to identify specific DNAm sites with extreme estimates of *S*. In addition, we considered DNAm sites associated with complex traits that have previously been shown to exhibit signatures of negative selection with *BayesS*^3^. DNAm sites associated with birthweight in EWAS had a higher proportion of DNAm sites with negative estimates of *S* compared to heritability matched DNAm sites.

For traits which are less polygenic it can be particularly hard to separate the actions of natural selection and genetic drift, which can generate extreme changes to the frequency of SNPs between human populations such as our study population (Europeans) and the common ancestor in which DNAm evolved (predating the out-of-Africa event)^7^. A Bayesian model accounting for genetic drift found that individual estimates of *S* could be due to either selection or genetic drift, but collectively DNAm was impacted by both positive and negative selection, not explainable by genetic drift alone^7^.

In addition to this, we were also able to characterize SNP based heritability and polygenicity for individual DNAm sites. Across all DNAm sites, average SNP based heritability was 28.6%. This is higher than previous estimates looking at ARIES data, but likely reflects the fact we are constrained to considering DNAm sites which pass convergence checks^52^. For each group of DNAm sites, average polygenicity was low. We found a striking relationship between polygenicity and selection, with positively selected DNAm associated with only a small number of mQTLs which together explained most of the heritability of the trait. Conversely, negatively selected DNAm is likely to be explained by more mQTLs, many of which we lack statistical power to identify. The genetic architecture of DNAm has been shown to have a large *cis*-mQTL effect plus polygenic *trans*-mQTLs of low effect sizes^19,52^. Studies in larger and more diverse populations should be undertaken to further investigate the relationship between polygenicity and selection. Our results provide insight into how genetic architecture of individual DNAm sites has been influenced by natural selection.

There are several limitations of this study. The model is restricted to looking at DNAm sites which pass MCMC convergence checks, which typically are those with high heritability in twin studies^12^. In addition, we compared *BayesNS* estimates to other selection metrics (*F_st_*, SDS, XPEHH, iHS), which are specialised to detect signatures of positive selection and have an estimate per SNP. This means that they do not make an ideal comparison group, since *BayesS* can be used to make inferences about both positive and negative selection and estimates are provided at the trait level. Blood cell counts have previously been reported to show signatures of negative selection^6^. Whilst our DNAm data has been adjusted for recorded cell counts^19^, the relationship between DNAm and blood cell counts^53^ could warrant further investigation in regards to whether it influences estimates of *S*. As mentioned, larger sample sizes are needed to detect mQTLs with low effect size which we are not powered to detect. Our study was also limited to the 450k array which measures 1.5% of the genome and is biased to promoters^29^. Large epidemiological studies profiled with EPIC arrays^54^ (measuring regulatory elements) are expected to find additional signatures for selection.

Overall, our study finds evidence for both positive and negative selection in the genetic architecture of DNAm. We are unable to cleanly place DNAm in the causal pathway between genetic variation and selection. Our results are consistent with two competing hypotheses; firstly, that selection occurs on DNAm due to a biological function it has, or secondly DNAm is influenced by complex traits that are themselves the target of selection. The presence of both positive and negative selection is an indication that both pathways may play a role. Specifically, we hypothesise that DNAm may perform a biological function which is of less selective importance than the complex traits that have widespread impact on genome wide DNAm, swamping and confusing the signal with a mixture of proximal and distal signals. Future work looking into the biological relevance of individual DNAm sites with positive and negative estimates of *S* could help to identify biological pathways which effect fitness. DNAm data from diverse individuals will be essential in separating the effects of drift and selection. Understanding the selective forces shaping DNAm could ultimately help identify potential targets for disease intervention.

## Methods

### Study Population

Participants were from the Avon Longitudinal Study of Parents and Children (ALSPAC)^55,56^, a large prospective cohort study that recruited 14,541 pregnancies, resident in the Bristol and Avon area with expected delivery dates between the 1^st^ of April 1991 and the 31^st^ of December 1992. Full details of the cohort have been published elsewhere^55,56^. The study website contains details of all the data that are available through a fully searchable data dictionary (http://www.bristol.ac.uk/alspac/researchers/our-data/). Written and informed consent has been obtained for all ALSPAC participants. Ethical approval for the study was obtained from the ALSPAC Ethics and Law Committee and the Local Research Ethics Committees (http://www.bristol.ac.uk/alspac/researchers/research-ethics/).

### ALSPAC genotype data

ALSPAC mothers were genotyped using the Illumina Human660W-quad array at Centre National de Génotypage (CNG). ALSPAC offspring were genotyped using the Illumina HumanHap550 quad chip genotyping platforms by 23andMe subcontracting the Wellcome Trust Sanger Institute, Cambridge, UK and the Laboratory Corporation of America, Burlington, NC, USA. For ALSPAC mothers, SNPs with a MAF of <1%, a call rate of <95%, or evidence for violations of Hardy–Weinberg equilibrium (*p* < 1 × 10^−6^) were removed. For ALSPAC offspring, SNPs with MAF of <1%, a call rate of <95% or evidence for violations of Hardy–Weinberg equilibrium (*p* < 5 × 10^−7^), were removed. Cryptic relatedness within mothers and within offspring was measured as proportion of identity by descent (IBD < 0.1). All individuals with non-European ancestry were removed. Imputation of ALSPAC genetic data was performed on a combined mother and child dataset using Impute2 against the 1000 Genomes Project Phase 1 Version 3 reference panel.

Linkage disequilibrium (LD) pruning was undertaken using PLINK^57^ using the following settings (r^2^=0.1, window size=50kb). SNPs residing within the Major Histocompatibility Complex (MHC) (chr6: 25Mb: 35Mb) and lactase (LCT) regions (chr2: 129Mb: 144Mb) were removed as they are known to be under high selective pressure (build 37). This left 474,939 SNPs available for analysis.

### DNA methylation data

In ALSPAC, blood from 1018 mother-child pairs were selected for analysis as part of ARIES^34^ (http://www.ariesepigenomics.org.uk/). Following DNA extraction, samples were bisulphite converted using the Zymo EZ DNA Methylation™ kit (Zymo, Irvine, CA, USA), and DNA methylation was measured using the Illumina Infinium HumanMethylation450 (HM450) BeadChip. ARIES consists of DNAm measures at five time points (three time points for children: birth, childhood, and adolescence; and two for mothers: during pregnancy and at middle age). We utilised data on a total of 1774 individuals from the adolescence and middle age time points with both DNAm and genotype data (http://www.ariesepigenomics.org.uk/).

DNAm was adjusted using the GoDMC pipeline (described elsewhere)^19^. Briefly, we adjusted for sex, age at measurement, batch variables, smoking and predicted cell counts. Genetic principal components (PCs), non-genetic DNAm PCs were also calculated using the GoDMC pipeline, and a genetic kinship matrix was fitted using GRAMMAR^58^. The residuals of these analyses were rank transformed to have a mean 0 and variance 1.

We selected the following DNAm sites for analyses:

1. 2000 DNAm sites with twin heritability estimates between 0.99 and 0.79^12^ (referred to as ‘highly heritable’ DNAm sites)
2. 1580 DNAm sites identified as having greater epigenetic similarity than can be explained genetically, so-called ‘epigenetic supersimilarity’ (ESS) DNAm sites^14^
3. 513 DNAm sites forming an epigenetic biomarker of aging, PhenoAge^32^, that is predictive of all-cause mortality
4. 243 DNAm sites associated with BMI (*p* < 1×10^−4^) in a discovery EWAS of 2707 individuals of European ancestry^30^. Results were obtained and downloaded from the EWAS catalog^49^
5. 893 DNAm sites associated with birthweight (*p* < 1×10^−4^) in a cord blood EWAS meta-analysis of 6023 individuals of European ancestry^31^. Results were obtained and downloaded from the EWAS catalog^49^

To serve as comparison groups, we additionally ran *BayesNS* on 513, 243 and 893 background DNAm sites matched on GC/CpG content and heritability to PhenoAge, BMI and birthweight associated sites respectively.

### *BayesNS* Analysis

*BayesS* is a Bayesian mixed linear model (MLM) method that can jointly estimate SNP-based heritability 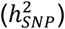, polygenicity (*π*) and the joint distribution between MAF and SNP effect size (*S*)^3^. The relationship between MAF and effect size is used to make inferences about natural selection and is modelled using the following mixture distribution as a prior for each SNP effect:

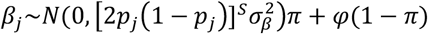

Where *β_j_* is the effect of a SNP *j*, *p_j_* is the MAF, 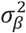 is the variance of SNP effects under a neutral model, *φ* is a point mass at zero and *π* is polygenicity (defined as the proportion of SNPs with non-zero effects). *S* is the estimated selection coefficient, when *S* > 0 effect size is positively related to MAF and when *S* < 0 effect size is negatively related to MAF. When *S* = 0, effect size and MAF are unrelated. *BayesS* uses a Markov Chain Monte Carlo (MCMC) algorithm for posterior inference. SNP-based heritability is estimated using the sampled effects of SNPs in the MCMC. We applied a nested version of *BayesS* (*BayesNS*) recommended for traits with low polygenicity such as DNAm. *BayesNS* considers SNPs together in non-overlapping windows and skips over regions of zero effect. SNPs in the same window are individually modelled as in *BayesS*, but also collectively considered as a window effect. The length of each window was set as 200kb, replicating the window size selected for analyses of gene expression^3^. Polygenicity (*π*) here is considered as the proportion of windows with nonzero effects. We considered each DNAm site as an individual trait in our analyses.

For the MCMC algorithm we set the chain length to 30,000 iterations with the first 10,000 discarded as burn-in. We plotted MCMC trace plots using *bayesplot* (http://mc-stan.org/bayesplot/) to visually assess convergence of the MCMC algorithm. In addition, we ran the Raftery and Lewis^35^ run length control diagnostic in *coda* and selected a threshold of less than 10 for the dependence factor (I) (Figures S2;S3). MCMC convergence checks were performed in R version 3.6.2.

### Accounting for Genetic Drift

We additionally ran a Bayesian model for genetic architecture which accounts for genetic drift. We used the MCMC algorithm from Ashraf and Lawson (2021)^7^ and applied it to the highly heritable, ESS and PhenoAge DNAm sites. Specifically, the prior for the selection coefficient *S*~*U*(−2,2), and for the standard deviation of *β* is *σ_β_*~*U*(0,2). Unlike in BayesS where *β_i_* is a prior, it and the allele frequency *pi* are treated as data, via the same relationship:

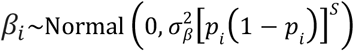

The *drift model* below is the appropriate model accounting for genetic drift. To furter check that our results are consistent with BayesS we report results for three models:

1. *Null model:* this extends the likelihood to account for *PIP*. *p*_i_ is considered fixed, and the likelihood from each SNP weighted by its inclusion probability *w_i_*, = *PIP*(*i*). The log-likelihood is 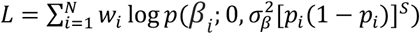, where *p* is the Normal distribution density.
2. *No-drift model:* no genetic drift but accounting for *PIP* and uncertainty in *β_i_*, *p*_i_ is considered fixed, the observed effect size 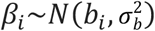 where *σ_b_* is the standard error of the estimate. Then following above 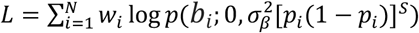.
3. *Drift model:* accounting for genetic drift, *PIP* and uncertainty in *β*_i_. Drift is modelled with the Baldings-Nichols model. Let *f_i_* be the true frequency in the ancestral population and *p_i_*, be observed as above. Then 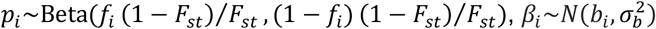 and 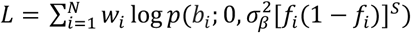.

*F_st_* is set to 0.15, matching the empirical estimate from the out-of-Africa bottleneck at these SNPs as in the original implementation^7^.

### Calculating variance explained

Following *BayesNS* analyses we investigated SNPs associated with DNAm at individual DNAm sites.

We selected SNPs with a high posterior inclusion probability (*PIP*) > = 0.8. We calculated the number of SNPs (nSNPs) with a *PIP* > = 0.8 for each DNAm site.

For each DNAm site we calculated variance explained for SNPs with *PIP* > = 0.8:

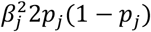

Where *β_j_* is the effect of a SNP *j*, *p_j_* is the MAF.

### Analysing LD score and selection metrics

To determine whether our results were influenced by LD, we additionally looked up European LD scores from the 1000 Genomes Project for each of these SNPs^43^. To compare *BayesNS* estimates of *S* with other selection scores we used metrics from the 1000 Genomes Selection Browser 1.0. We selected the same five annotations used in GoDMC^19^, reflecting selection over different timescales: singleton density score (SDS^17^; UK10K), *F_st_* ^40^(Global *F_st_* (CEU vs. YRI vs. CHB)), integrated haplotype score (iHS;CEU)^41^, cross population extended haplotype homozygosity (XPEHH; CEU vs. YRI) and XPEHH (CEU vs. CHB)^42^. These methods focus on positive selection^18^; *F_st_* is based on population differentiation^40^, XPEHH is a cross-population test based on extended haplotype homozygosity (EHH), iHS is defined as the log ratio of integrated haplotype scores for each allele in a single population^41^. SDS measures very recent changes in allele frequency from contemporary genome sequences and has been applied to the UK10K dataset^17^.

For each DNAm site we calculated the mean value for each of these selection scores for SNPs with *PIP* > 0.1 and *PIP* > 0.001 respectively. Each mean was weighted by the SNPs *PIP* value. We used *ggpairs* in R version 3.6.2 to plot pairwise distributions of *BayesNS,* LDSC and selection scores and to compute the Pearson correlation coefficient between these variables.

### Enrichment Analysis

We assessed enrichment or depletion of DNAm sites for 25 chromatin states and TFBSs in 127 different cell types comprising 30 tissues. These data were generated by the Roadmap Epigenomics Project^44^ (http://www.roadmapepigenomics.org/) and ENCODE (https://www.encodeproject.org/). We used Locus Overlap Analysis (LOLA)^59^ (Bioconductor version: Release 3.12) to perform a two-sided Fisher’s exact test. Since the magnitude of *S* reflects the strength of selection we selected DNAm sites with estimates of *S* ≥ 0.5 and *S* ≤ −0.5 for analyses. Background sites from the HumanMethylation450 array were matched on GC and CpG content and heritability prior to analysis (Figure S6), as differential GC content/heritability’s between the sites of interest and background sites may bias the results. Analyses were conducted using R v. 3.6.2.

Four groups of DNAm sites were considered for enrichment analysis:

1. 212 highly heritable DNAm sites with estimates of *S* ≥ 0.5 (*positive highly heritable DNAm sites*)
2. 376 highly heritable DNAm sites with estimates of *S* ≤ −0.5 (*negative highly heritable DNAm sites*)
3. 123 ESS DNAm sites with estimates of *S* ≥ 0.5 (*positive ESS DNAm sites*)
4. 218 ESS DNAm sites with estimates of *S* ≤ −0.5 (*negative ESS DNAm sites*)

### DNAm and complex traits

We ran *BayesNS* on DNAm sites associated with BMI and birthweight in two large-scale EWAS of participants with European ancestry. In addition, we ran *BayesNS* on DNAm sites not associated with the traits of interest, matched on GC/CpG content and heritability. We split DNAm sites in each set (PhenoAge, BMI-associated DNAm sites, birthweight-associated DNAm sites and matched background DNAm sites) into two groups: DNAm sites with negative estimates of *S* ≤ −0.5 and DNAm sites with estimates of *S* > −0.5. We then performed one-sided Fisher’s exact tests to investigate whether DNAm sites associated with PhenoAge, BMI and birthweight exhibit statistically different estimates of S compared to a set of matched background DNAm sites. We additionally performed GOterm enrichment analysis implemented in missmethyl^60,61^.

## Acknowledgements

We are extremely grateful to all the families who took part in this study, the midwives for their help in recruiting them, and the whole ALSPAC team, which includes interviewers, computer and laboratory technicians, clerical workers, research scientists, volunteers, managers, receptionists, and nurses.

## Author contributions

Analysed the data: C.H., G.H., J.L.M., D.J.L., T.R.G., S.R.

Contributed data: GoDMC, ALSPAC

Designed and managed the study: J.L.M., D.J.L., T.R.G., S.R.

Wrote the manuscript: C.H., J.L.M., D.J.L., T.R.G., S.R.

