## Supplementary Figures for "Evidence of positive and negative selection associated with DNA methylation"

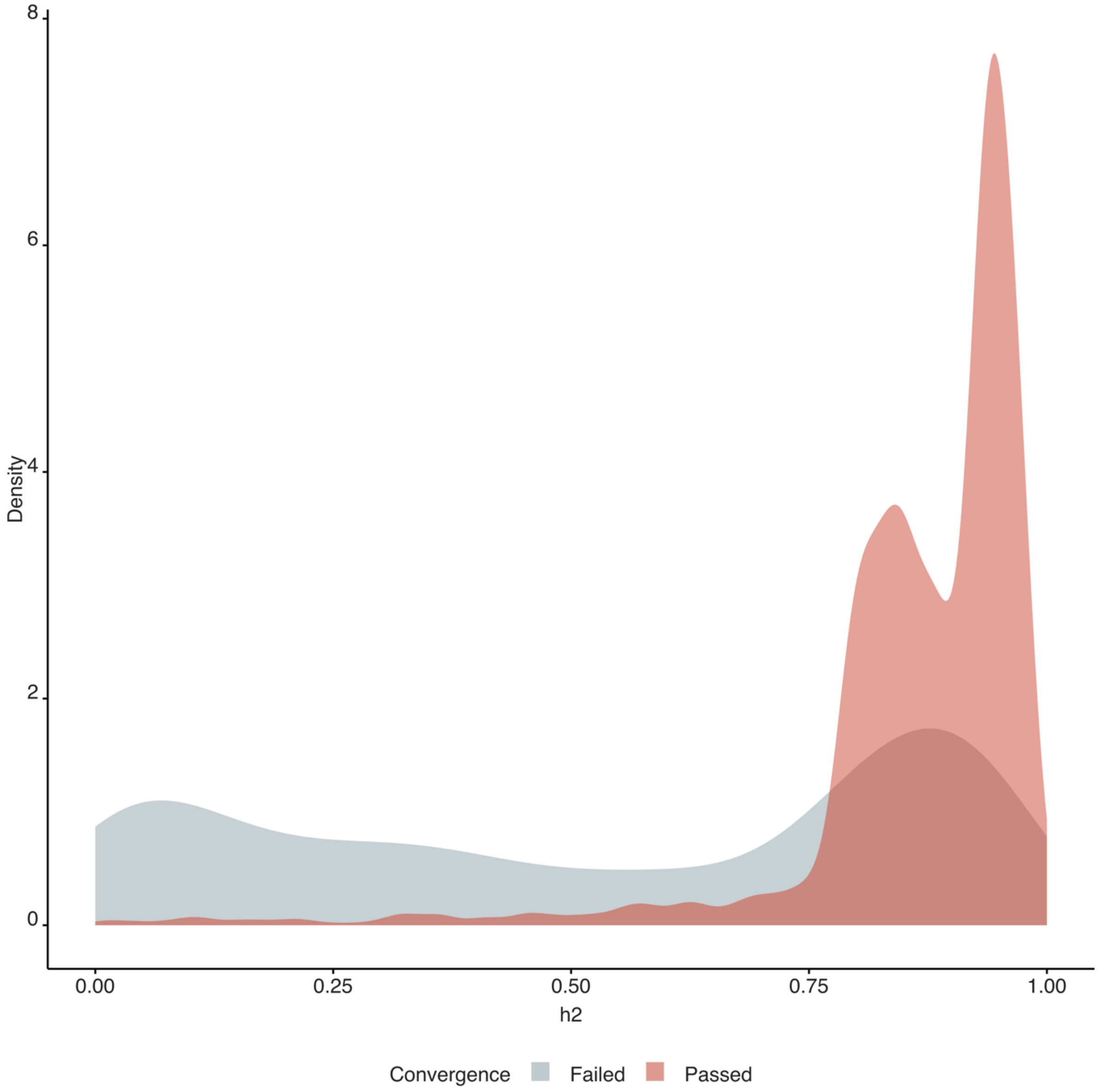


**Figure S1 |** Density plot of twin heritability estimates[1] across all DNAm sites (highly heritable, ESS and PhenoAge) coloured by whether they failed (grey) or passed (red) MCMC convergence tests.

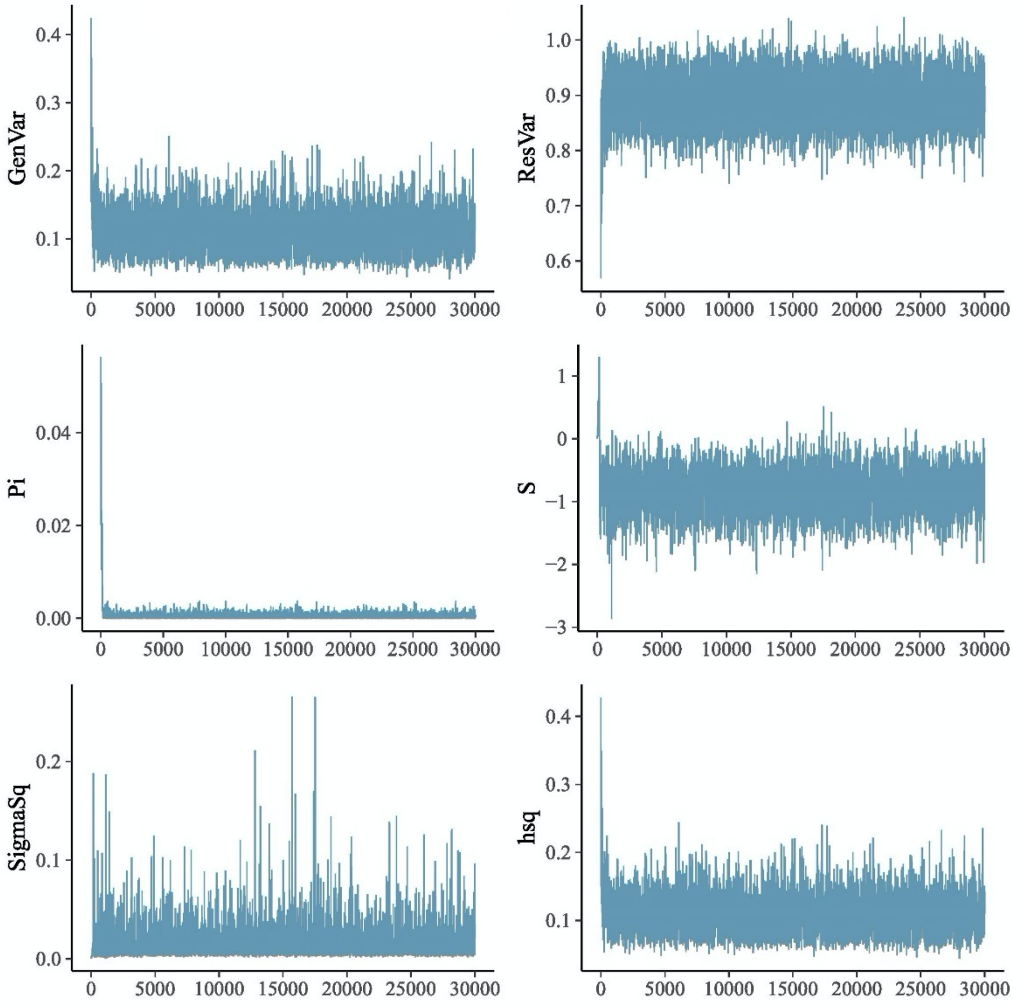


**Figure S2 A|** Example of a DNAm site which passed visual convergence checks. Bayesplot of MCMC algorithm.

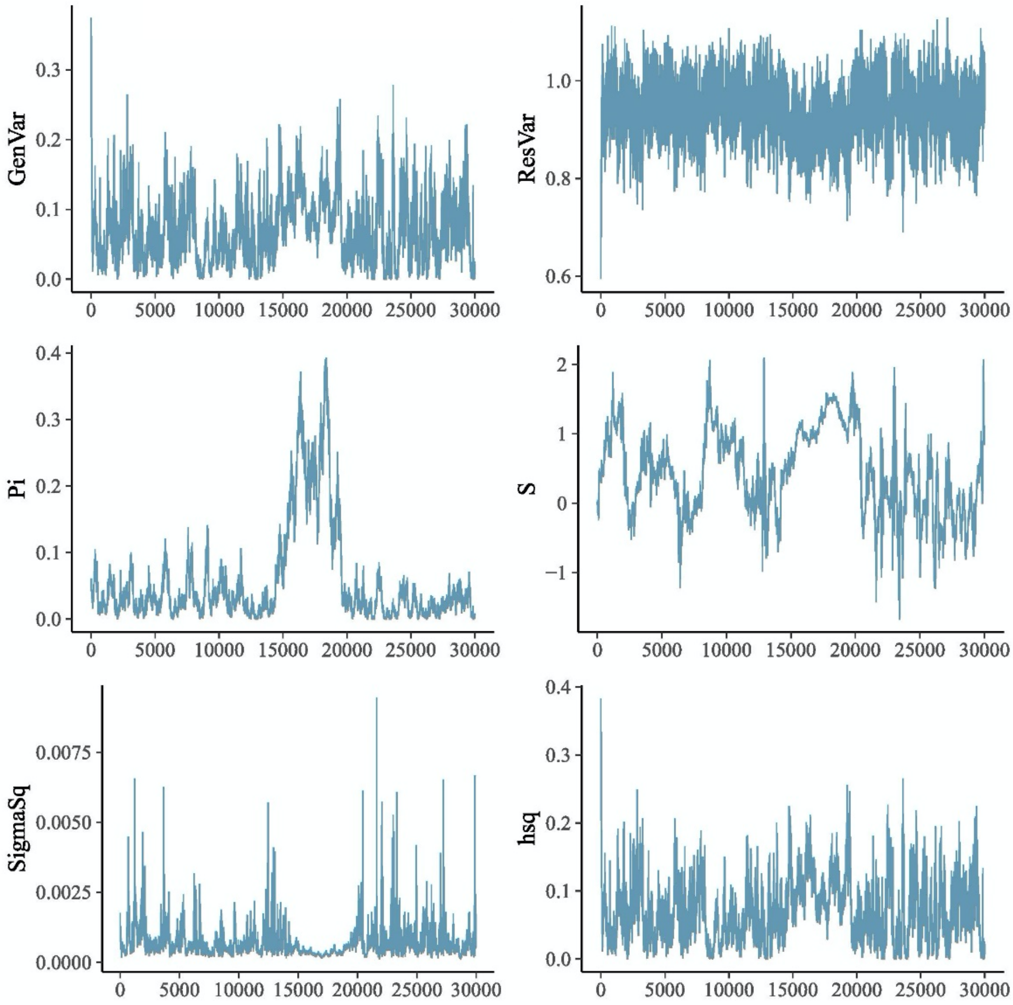


**Figure S2 B |** Example of a DNAm site which failed visual convergence checks. Bayesplot of MCMC algorithm.

### Highly Heritable Probes

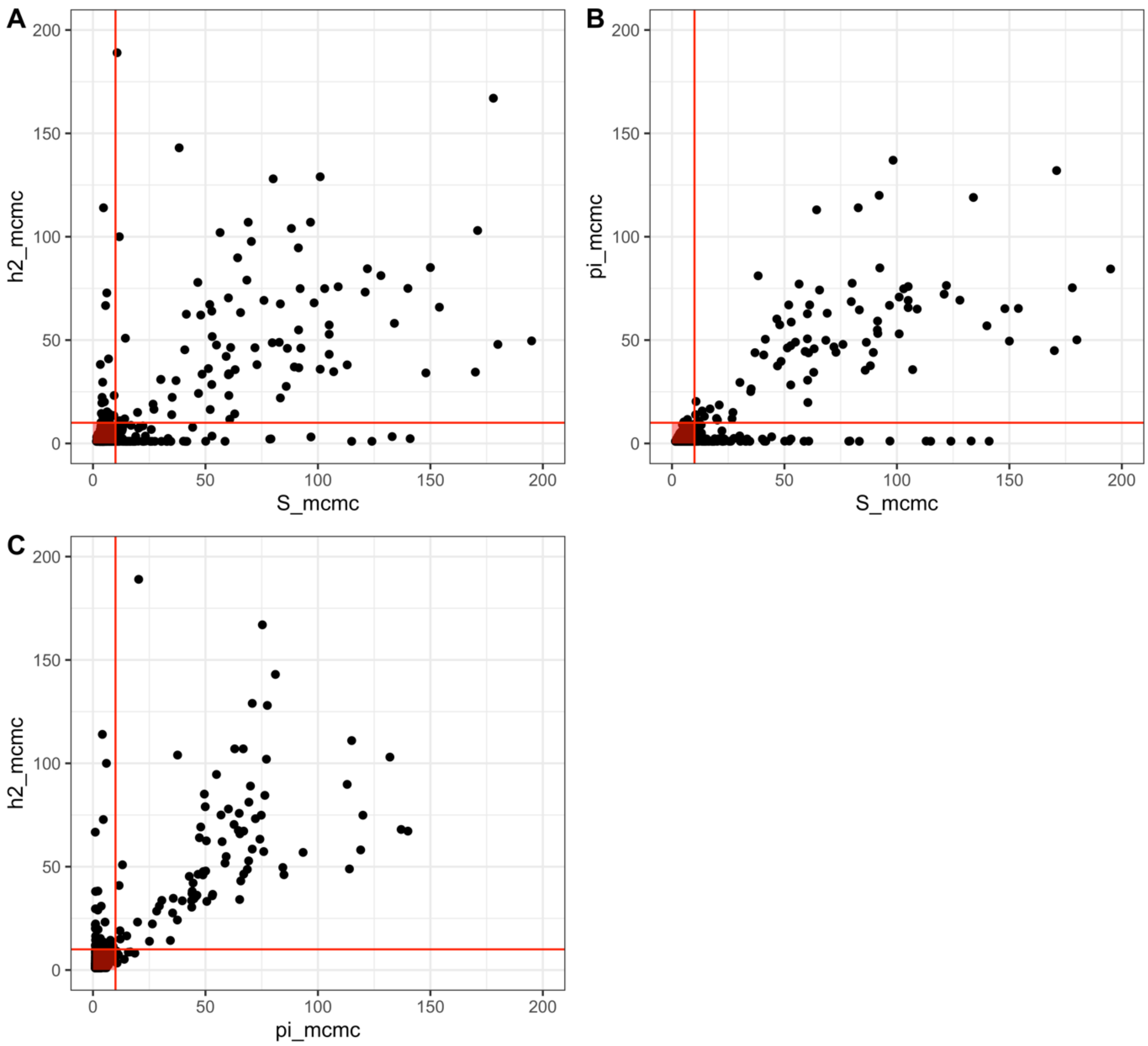


**Figure S3.1 |** Dependence Factor (I) from Raferty-Lewis[3] length control diagnostic for the MCMC algorithm for the highly heritable DNAm sites. Threshold of 10 was selected for I (shown by red lines). Probes taken forward had an I value of less than 10 for *S*, $h^2_{SNP}$ and pi ($\pi$).

### ESS Probes

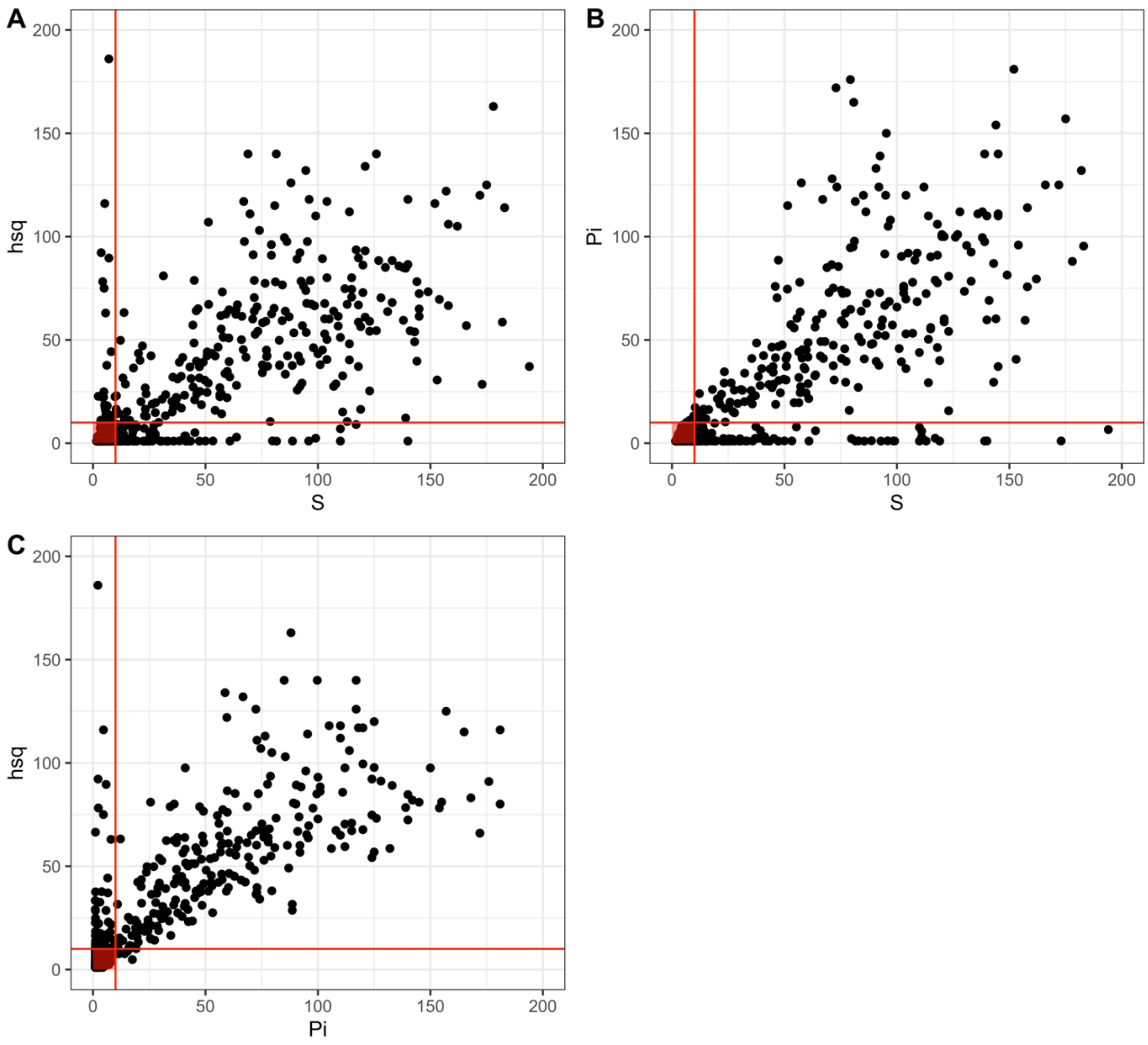


**Figure S3.2|** Dependence Factor (I) from Raferty-Lewis[3] length control diagnostic for the MCMC algorithm for the ESS DNAm sites. Threshold of 10 was selected for I (shown by red lines). Probes taken forward had an I value of less than 10 for *S*, $h^2_{SNP}$ and pi ($\pi$).

**PhenoAge Probes**

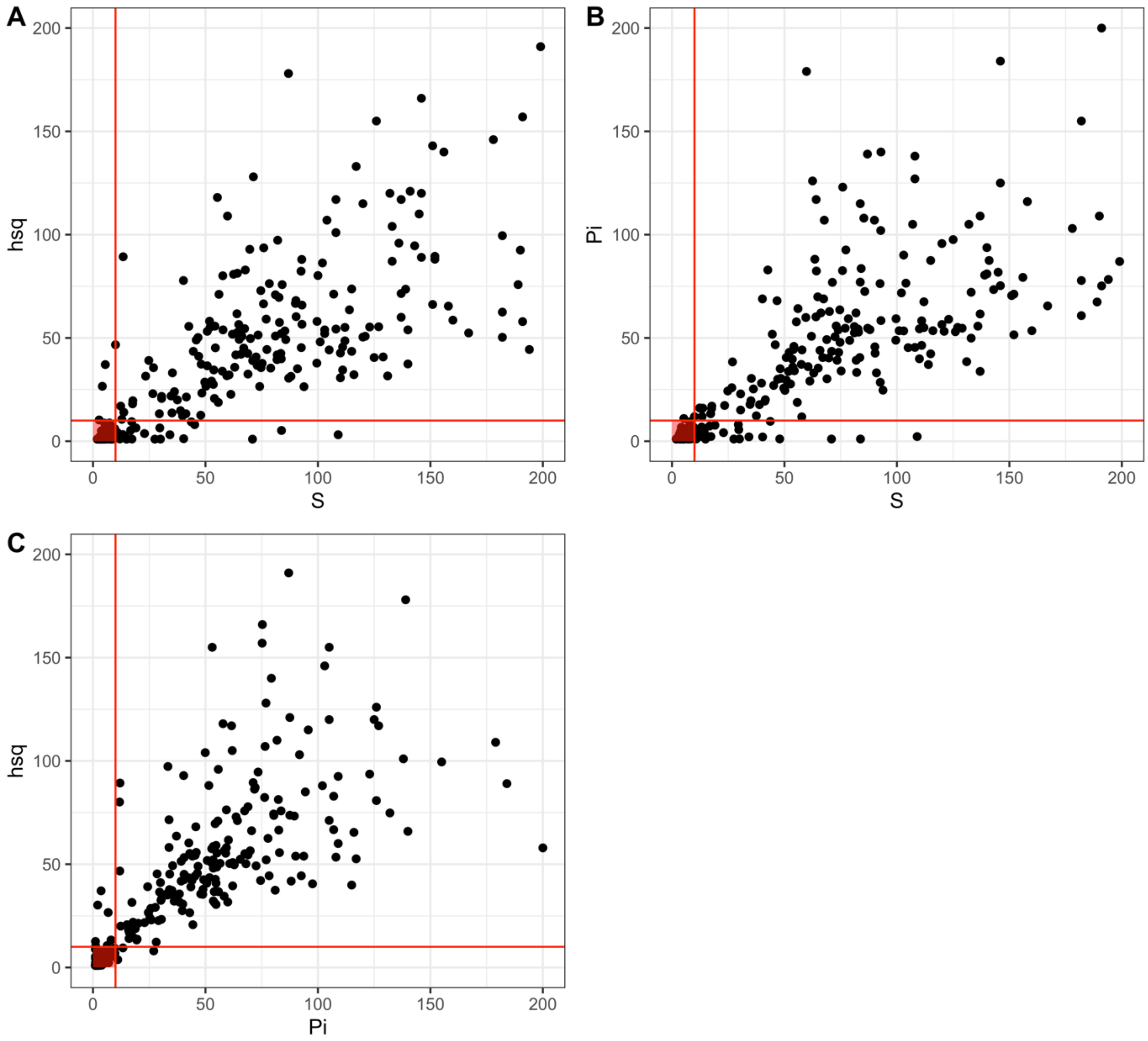


**Figure S3.3 |** Dependence Factor (I) from Raferty-Lewis[3] length control diagnostic for the MCMC algorithm for the PhenoAge DNAm sites. Threshold of 10 was selected for I (shown by red lines). Probes taken forward had an I value of less than 10 for *S*, $h^2_{SNP}$ and pi ($\pi$).

**BMI Probes**

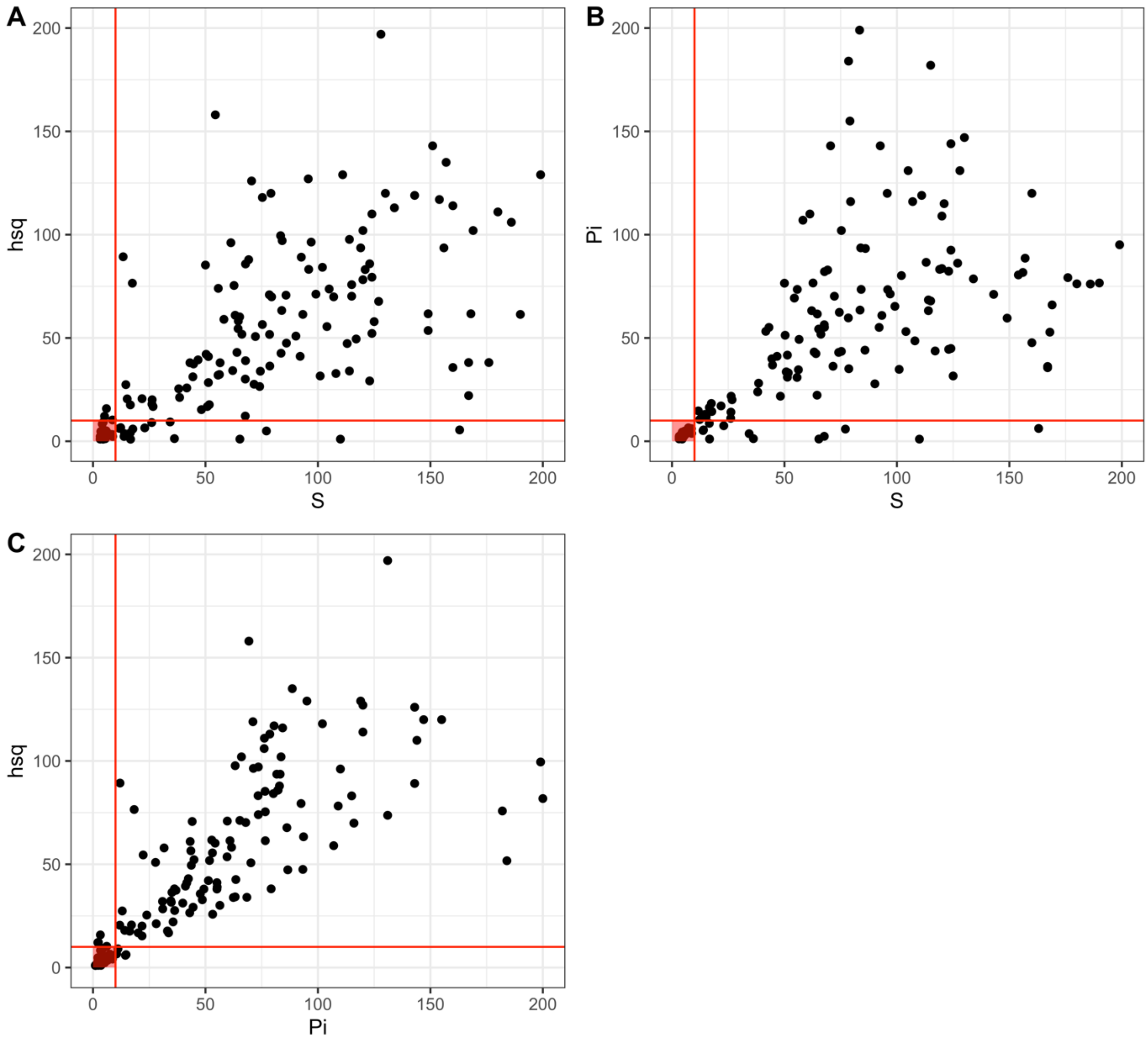


**Figure S3.3 |** Dependence Factor (I) from Raferty-Lewis[3] length control diagnostic for the MCMC algorithm for the BMI-associated DNAm sites. Threshold of 10 was selected for I (shown by red lines). Probes taken forward had an I value of less than 10 for *S*, $h^2_{SNP}$ and pi ($\pi$).

**Birthweight Probes**

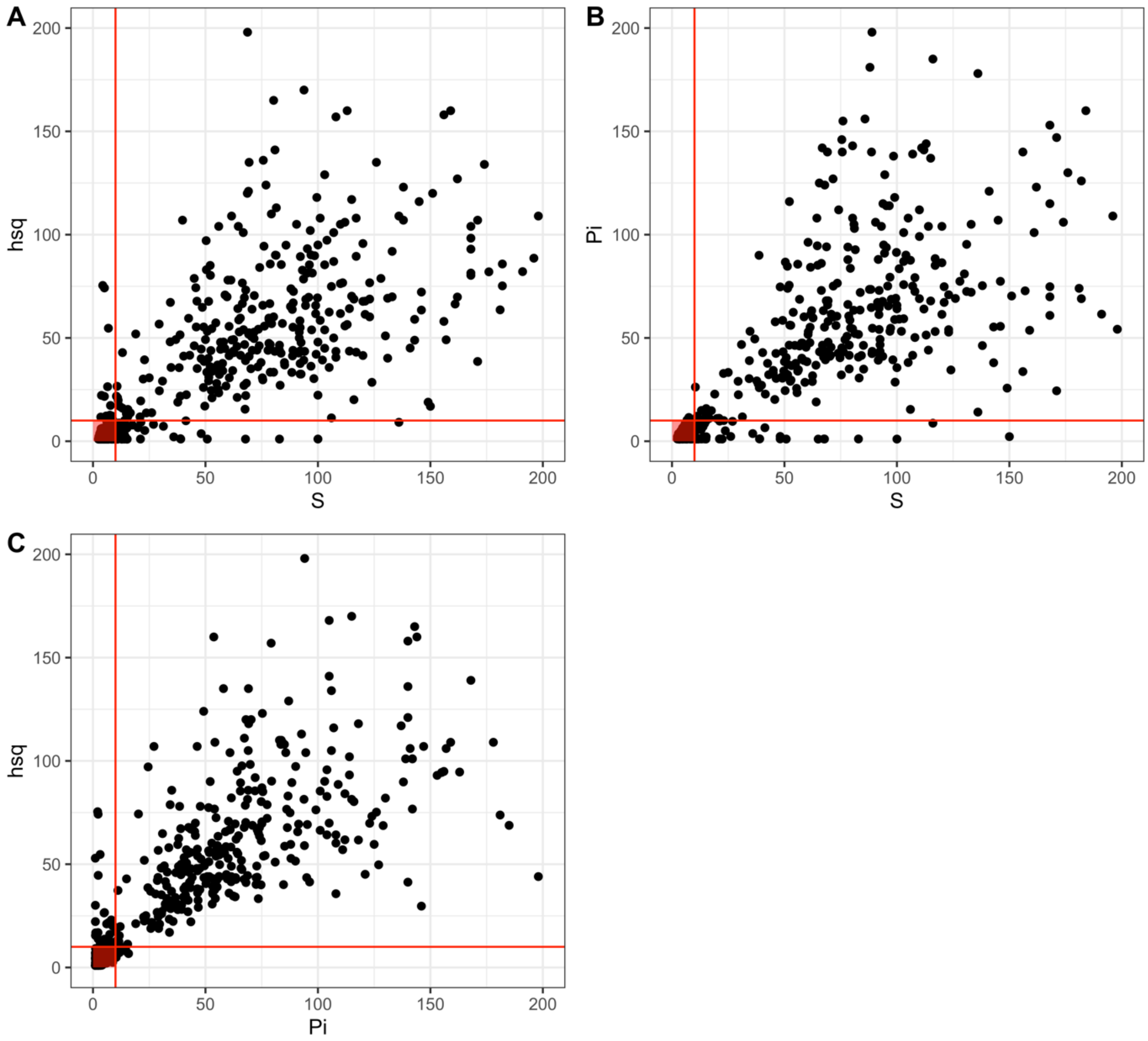


**Figure S3.4 |** Dependence Factor (I) from Raferty-Lewis[3] length control diagnostic for the MCMC algorithm for the birthweight-associated DNAm sites. Threshold of 10 was selected for I (shown by red lines). Probes taken forward had an I value of less than 10 for *S*, $h^2_{SNP}$ and pi ($\pi$).

### Highly Heritable DNAm sites

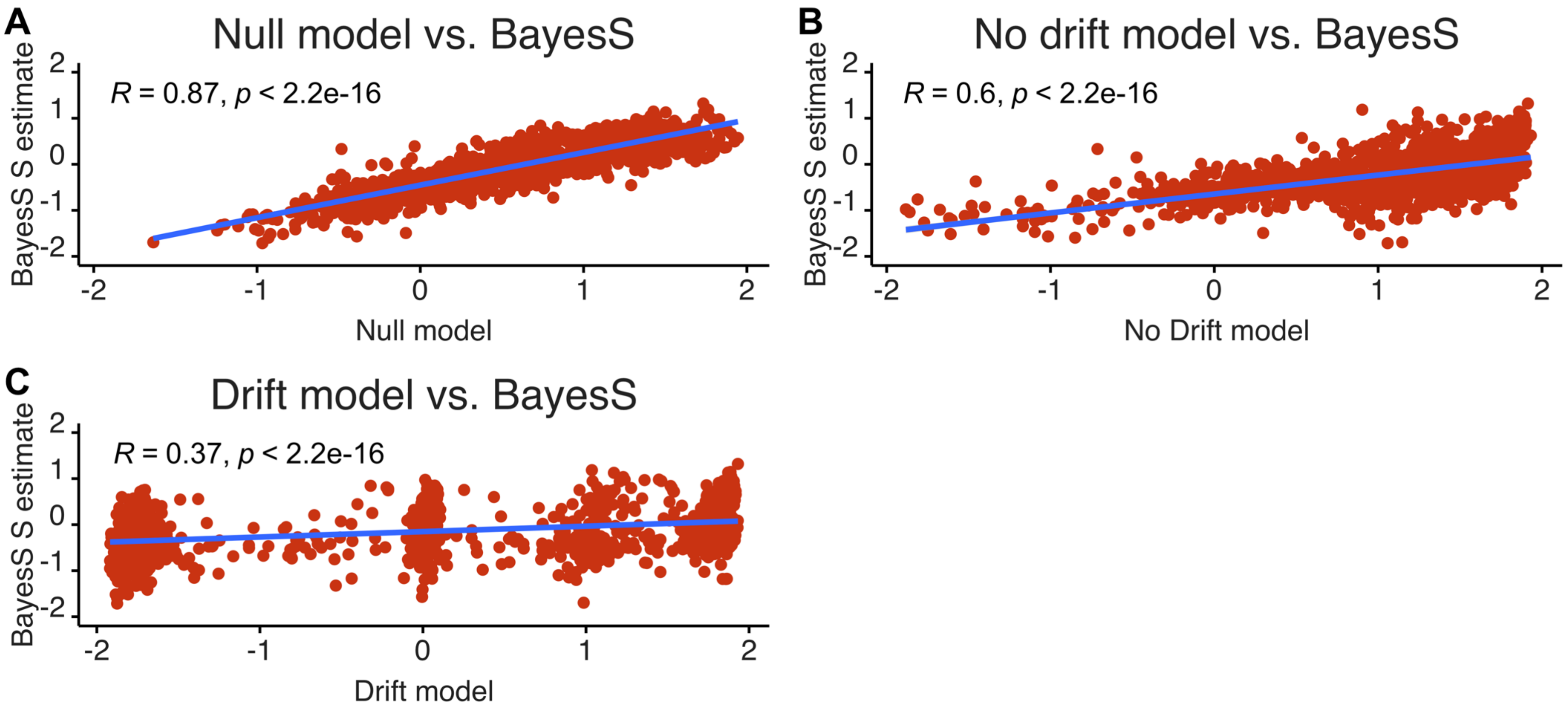


**Figure S4.1 | Comparison of *BayesNS* estimates of *S* with model correcting for genetic drift[4] for 1804 highly heritable DNAm sites.** Estimates of *S* from *BayesNS* plotted against estimates from **(A)** *null model:* an implementation of *BayesNS* which accounts for posterior inclusion probability (PIP) (R=0.87), **(B)** *no-drift model:* no genetic drift but accounting for PIP and uncertainty in betas (R=0.60), **(C)** d*rift model:* accounting for genetic drift, PIP and uncertainty in betas (R=0.37).

### ESS DNAm sites

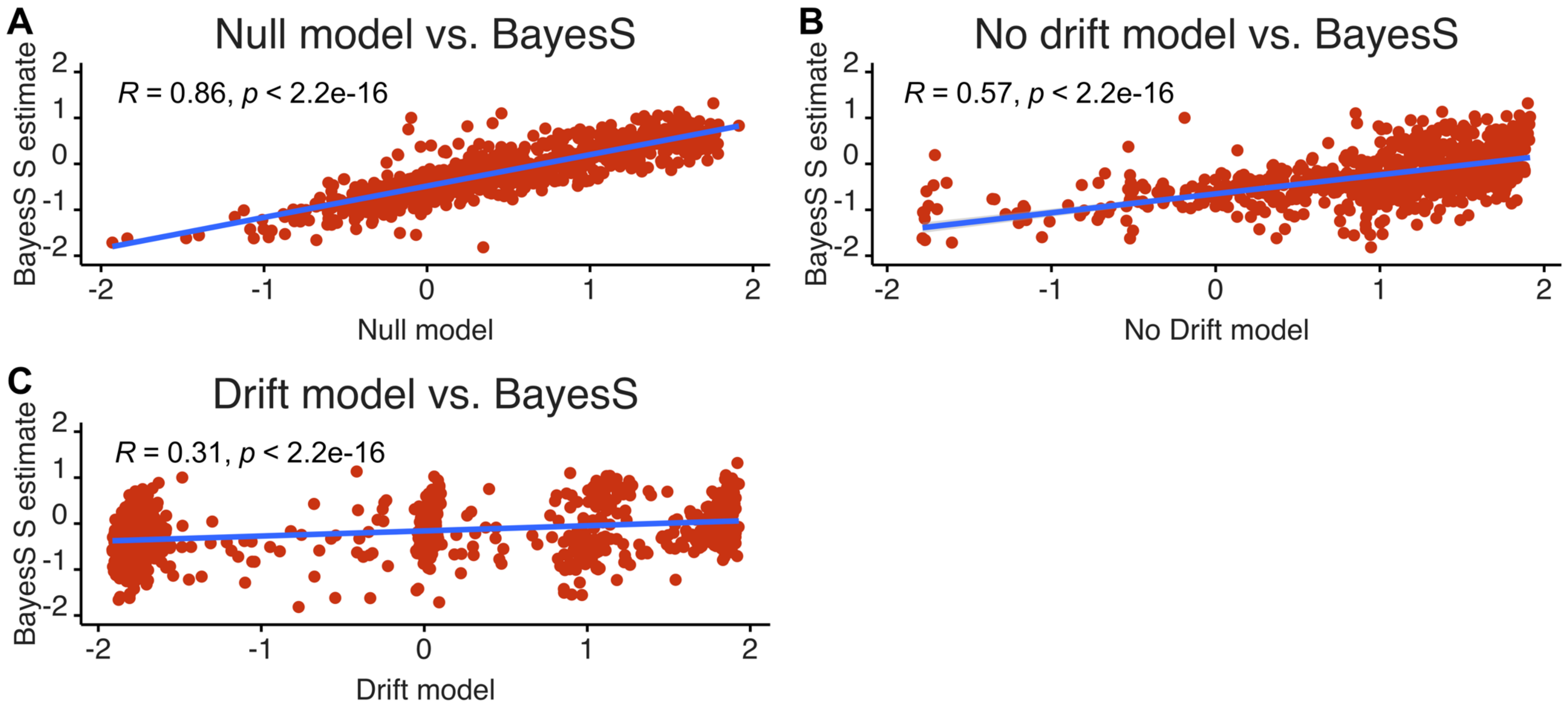


**Figure S4.2 | Comparison of *BayesNS* estimates of *S* with model correcting for genetic drift[4] for 887 ESS DNAm sites.** Estimates of *S* from *BayesNS* plotted against estimates from **(A)** *null model:* an implementation of *BayesNS* which accounts for posterior inclusion probability (PIP) (R=0.86), **(B)** *no-drift model:* no genetic drift but accounting for PIP and uncertainty in betas (R=0.57), **(C)** d*rift model:* accounting for genetic drift, PIP and uncertainty in betas (R=0.31).

### PhenoAge DNAm sites

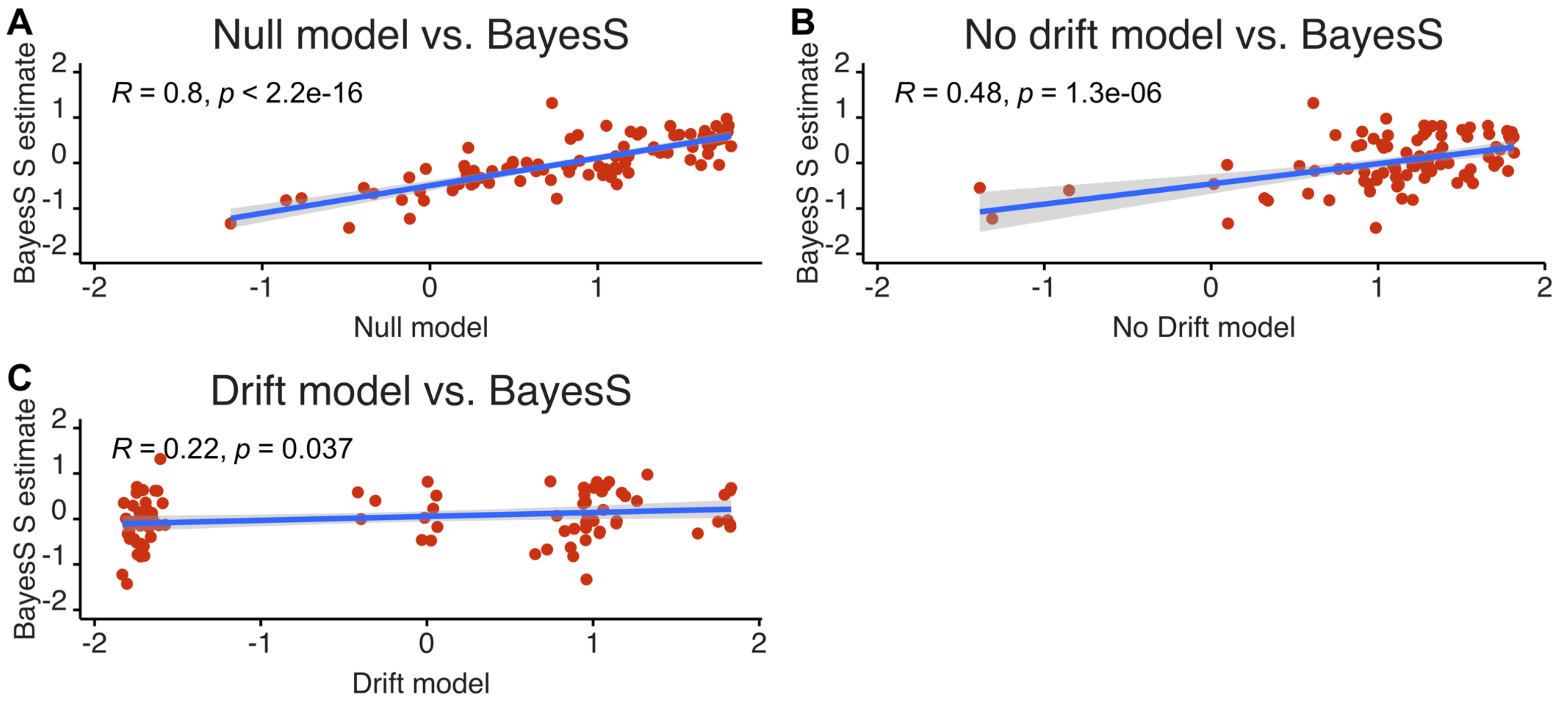


**Figure S4.3 | Comparison of *BayesNS* estimates of *S* with model correcting for genetic drift[4] for 74 PhenoAge DNAm sites.** Estimates of *S* from *BayesNS* plotted against estimates from **(A)** *null model:* an implementation of *BayesNS* which accounts for posterior inclusion probability (PIP) (R=0.80), **(B)** *no-drift model:* no genetic drift but accounting for PIP and uncertainty in betas (R=0.48), **(C)** d*rift model:* accounting for genetic drift, PIP and uncertainty in betas (R=0.22)

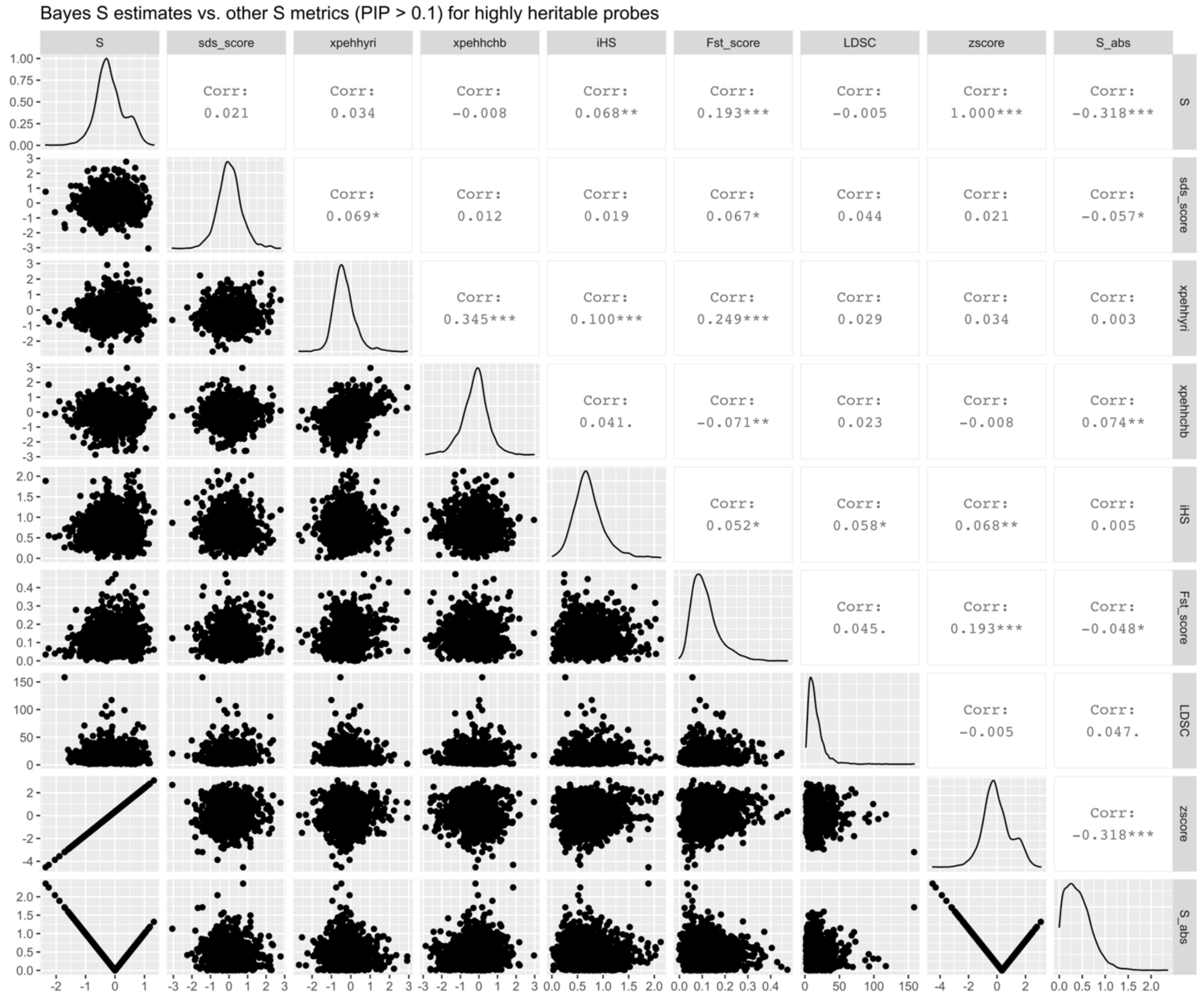


**Figure S5A| Correlations between *BayesNS* estimate of *S* and mean SDS, XPEHH-YRI, XPEHH-CHB, iHS, $F_{st}$ and LDSC weighted by PIP for SNPs with PIP > 0.1 for highly heritable probes[5,6].** Absolute value of *S* (S_abs) and z-score of *S* also shown. ******* indicates $p < 0.001$, ****** indicates $p < 0.01$, ***** indicates $p < 0.05$, **.** indicates $p < 0.1$. Figure generated using *ggpairs* in R v. 3.6.2. Abbreviations: *SDS*: singleton density score, *XPEHH- YRI* **(**cross population extended haplotype homozygosity (XPEHH; CEU vs. YRI), *XPEHH-CHB*: (CEU vs. CHB), *iHS*: integrated haplotype score, *LDSC*: Linkage disequilibrium score, *PIP*: posterior inclusion probability, *SNP*: single nucleotide polymorphism.

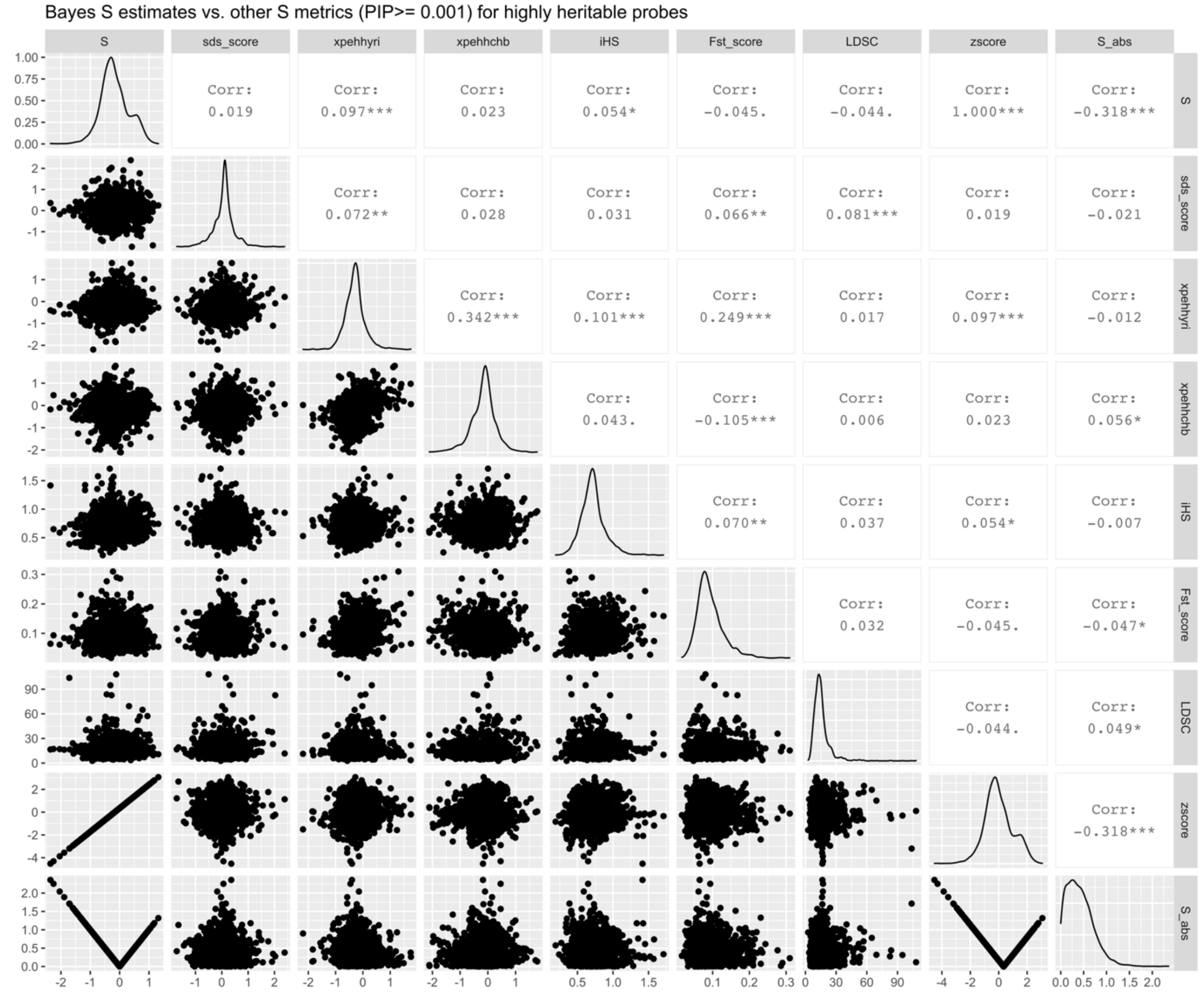


**Figure S5B| Correlations between *BayesNS* estimate of *S* and mean SDS, XPEHH-YRI, XPEHH-CHB, iHS, $F_{st}$ and LDSC weighted by PIP for SNPs with PIP >=0.001 for highly heritable probes[5,6].** Absolute value of *S* (S_abs) and z-score of *S* also shown. ******* indicates $p < 0.001$, ****** indicates $p < 0.01$, ***** indicates $p < 0.05$, **.** indicates $p < 0.1$. Figure generated using *ggpairs* in R v. 3.6.2. Abbreviations: *SDS*: singleton density score, *XPEHH- YRI* **(**cross population extended haplotype homozygosity (XPEHH; CEU vs. YRI), *XPEHH-CHB*: (CEU vs. CHB), *iHS*: integrated haplotype score, *LDSC*: Linkage disequilibrium score, *PIP*: posterior inclusion probability, *SNP*: single nucleotide polymorphism

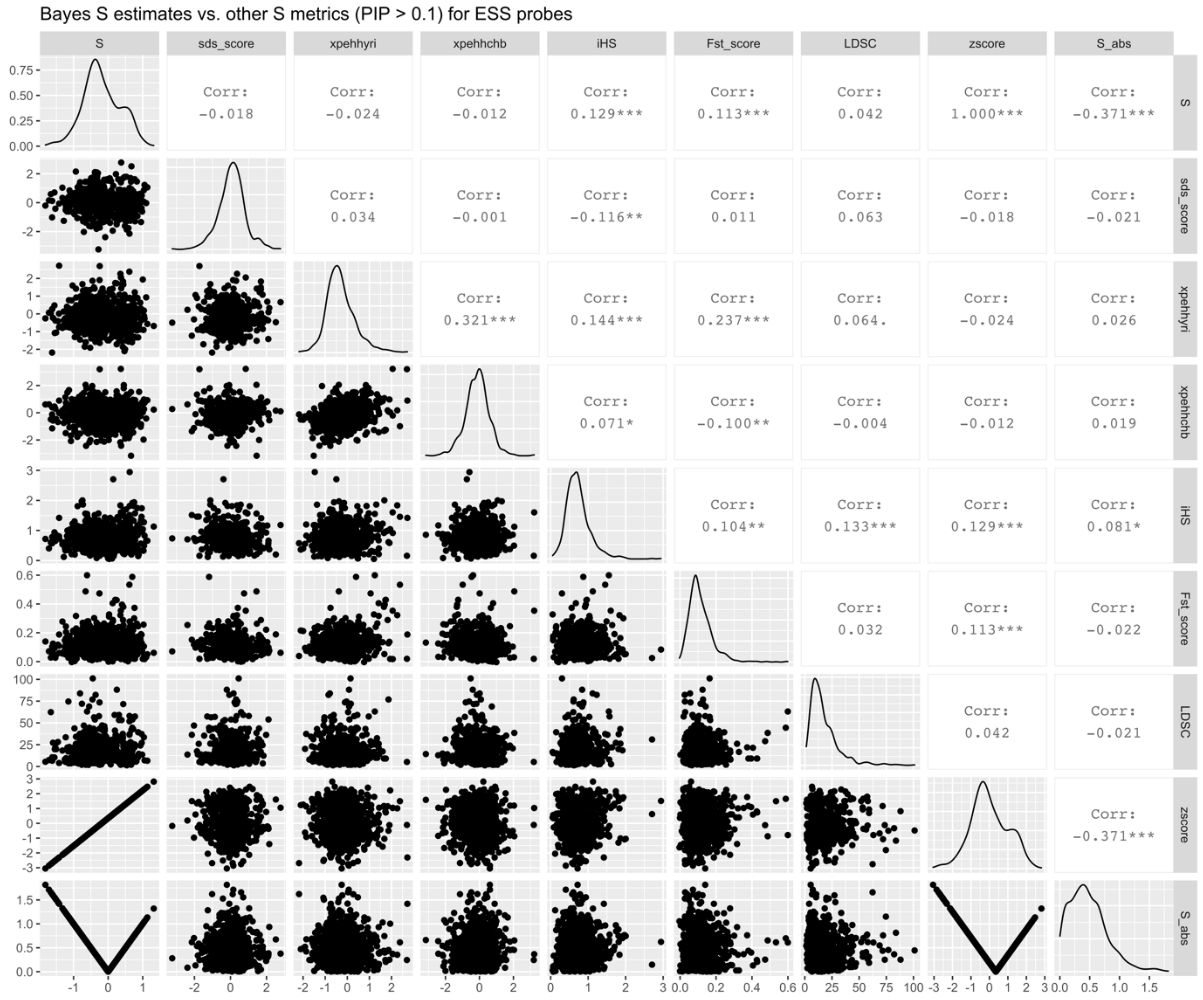


**Figure S5C | Correlations between *BayesNS* estimate of *S* and mean SDS, XPEHH-YRI, XPEHH-CHB, iHS, $F_{st}$ and LDSC weighted by PIP for SNPs with PIP > 0.1 for ESS probes[5,6].** Absolute value of *S* (S_abs) and z-score of *S* also shown. ******* indicates $p < 0.001$, ****** indicates $p < 0.01$, ***** indicates $p < 0.05$, **.** indicates $p < 0.1$. Figure generated using *ggpairs* in R v. 3.6.2. Abbreviations: *SDS*: singleton density score, *XPEHH- YRI* **(**cross population extended haplotype homozygosity (XPEHH; CEU vs. YRI), *XPEHH-CHB*: (CEU vs. CHB), *iHS*: integrated haplotype score, *LDSC*: Linkage disequilibrium score, *PIP*: posterior inclusion probability, *SNP*: single nucleotide polymorphism

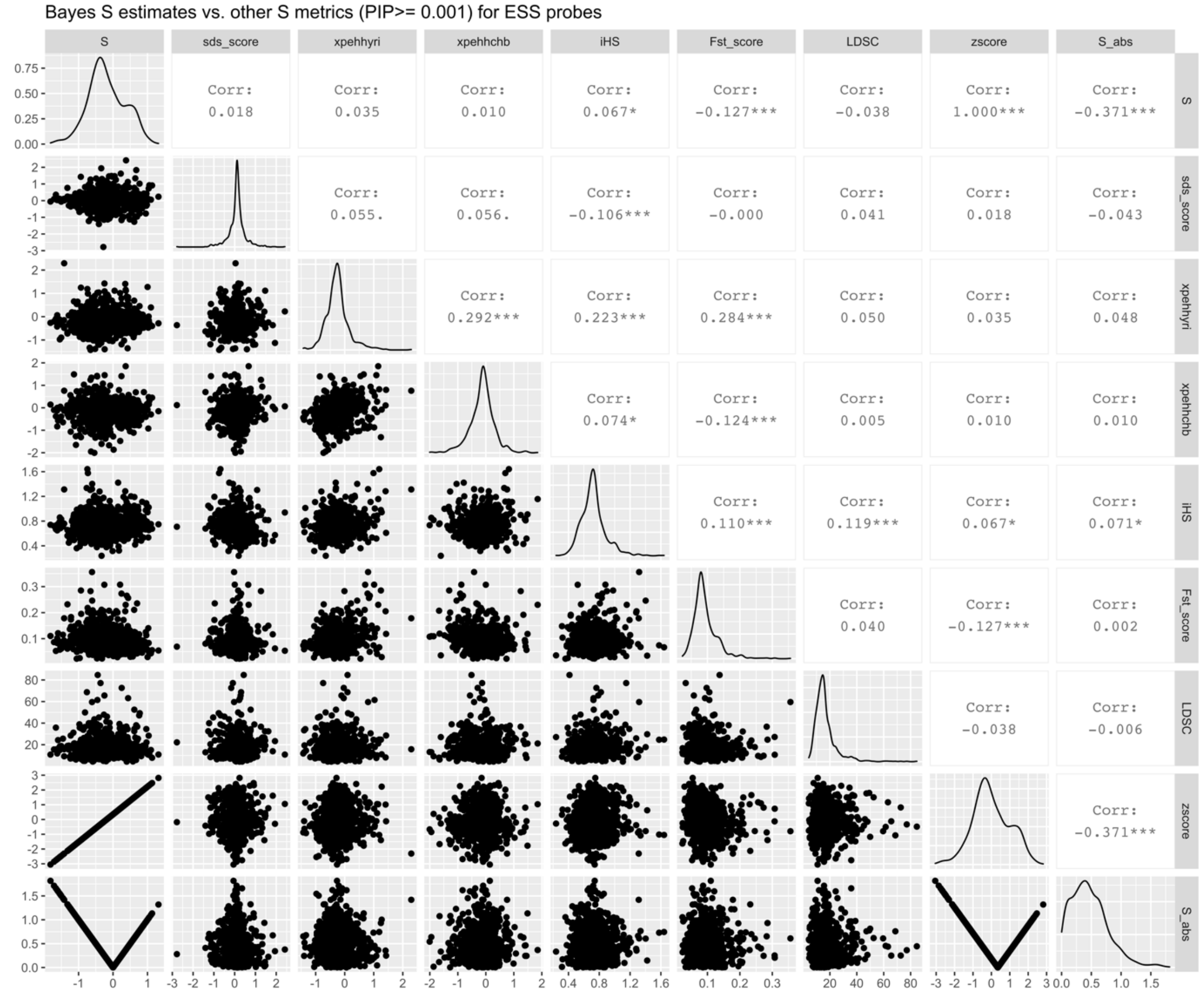


**Figure S5D | Correlations between *BayesNS* estimate of *S* and mean SDS, XPEHH-YRI, XPEHH-CHB, iHS, $F_{st}$ and LDSC weighted by PIP for SNPs with PIP >= 0.001 for ESS probes[5,6].** Absolute value of *S* (S_abs) and z-score of *S* also shown. ******* indicates $p < 0.001$, ****** indicates $p < 0.01$, ***** indicates $p < 0.05$, **.** indicates $p < 0.1$. Figure generated using *ggpairs* in R v. 3.6.2. Abbreviations: *SDS*: singleton density score, *XPEHH- YRI* **(**cross population extended haplotype homozygosity (XPEHH; CEU vs. YRI), *XPEHH-CHB*: (CEU vs. CHB), *iHS*: integrated haplotype score, *LDSC*: Linkage disequilibrium score, *PIP*: posterior inclusion probability, *SNP*: single nucleotide polymorphism

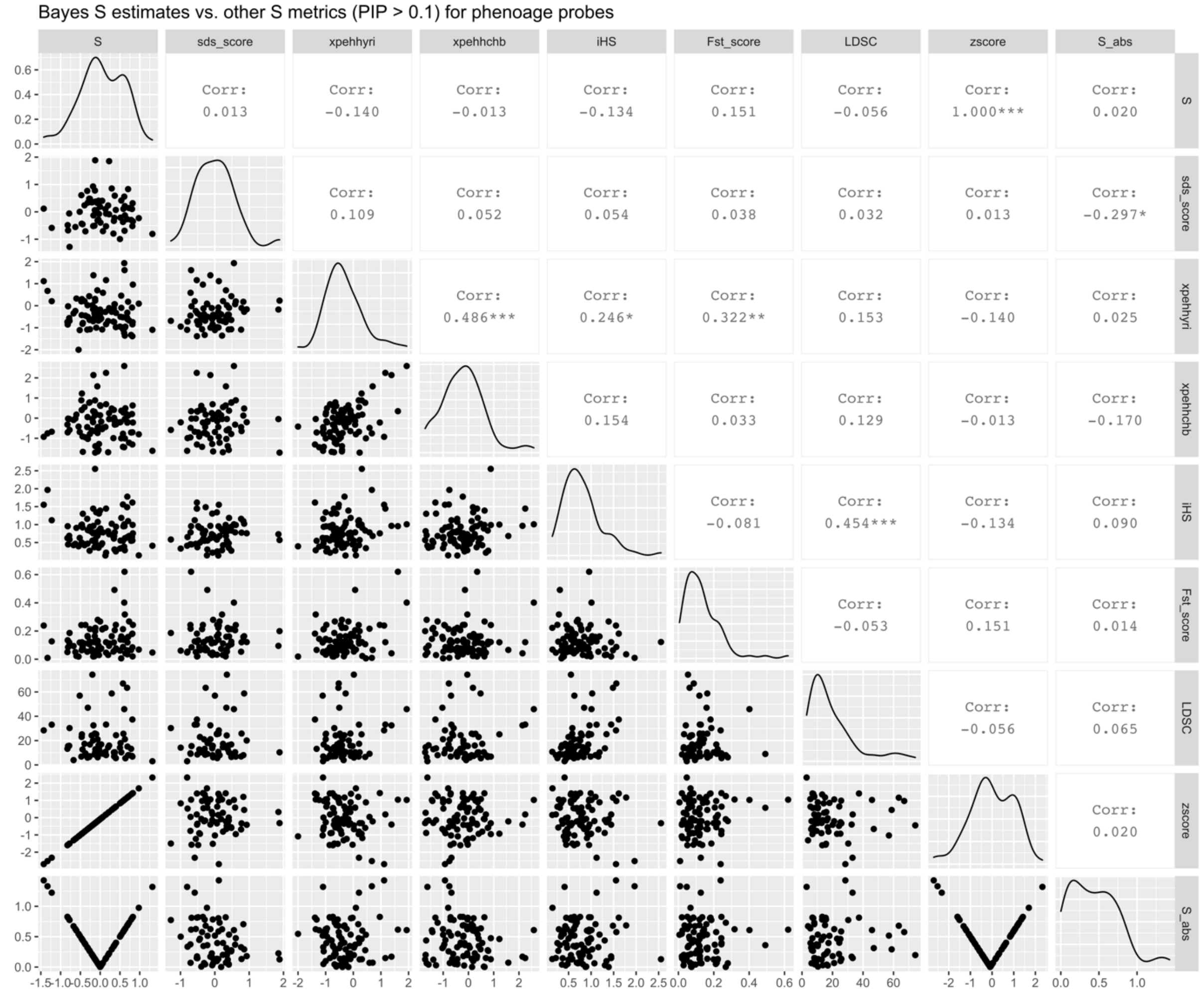


**Figure S5E | Correlations between *BayesNS* estimate of *S* and mean SDS, XPEHH-YRI, XPEHH-CHB, iHS, $F_{st}$ and LDSC weighted by PIP for SNPs with PIP > 0.1 for PhenoAge probes[5,6].** Absolute value of *S* (S_abs) and z-score of *S* also shown. ******* indicates *p* < 0.001, ****** indicates *p* < 0.01, ***** indicates *p* < 0.05, **.** indicates *p* < 0.1. Figure generated using *ggpairs* in R v. 3.6.2. Abbreviations: *SDS*: singleton density score, *XPEHH- YRI* **(**cross population extended haplotype homozygosity (XPEHH; CEU vs. YRI), *XPEHH-CHB*: (CEU vs. CHB), *iHS*: integrated haplotype score, *LDSC*: Linkage disequilibrium score, *PIP*: posterior inclusion probability, *SNP*: single nucleotide polymorphism

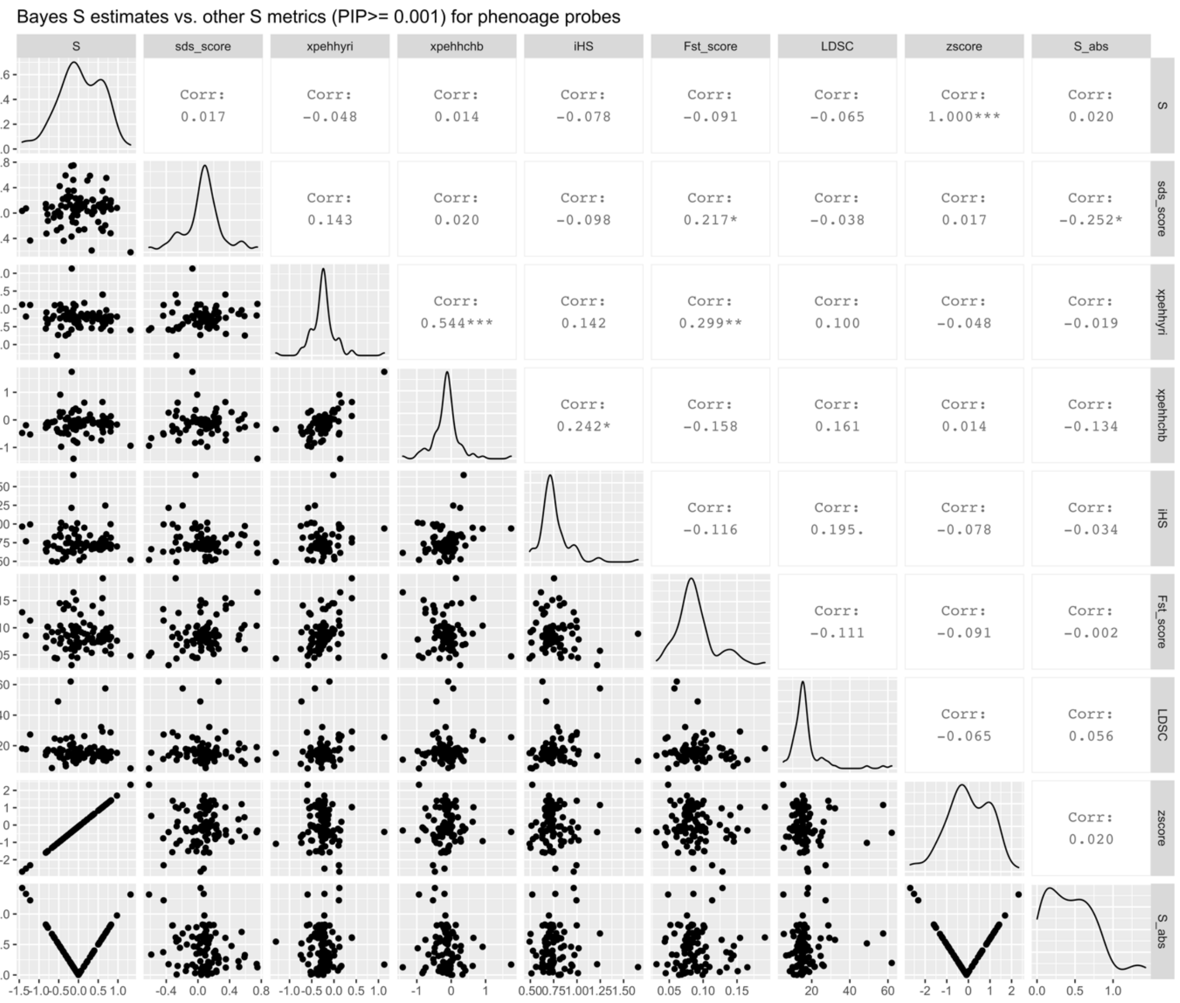


**Figure S5F | Correlations between *BayesNS* estimate of *S* and mean SDS, XPEHH-YRI, XPEHH-CHB, iHS, $F_{st}$ and LDSC weighted by PIP for SNPs with PIP >= 0.001 for PhenoAge probes[5,6].** Absolute value of *S* (S_abs) and z-score of *S* also shown. ******* indicates $p < 0.001$, ****** indicates $p < 0.01$, ***** indicates $p < 0.05$, **.** indicates $p < 0.1$. Figure generated using *ggpairs* in R v. 3.6.2. Abbreviations: *SDS*: singleton density score, *XPEHH- YRI* **(**cross population extended haplotype homozygosity (XPEHH; CEU vs. YRI), *XPEHH-CHB*: (CEU vs. CHB), *iHS*: integrated haplotype score, *LDSC*: Linkage disequilibrium score, *PIP*: posterior inclusion probability, *SNP*: single nucleotide polymorphism

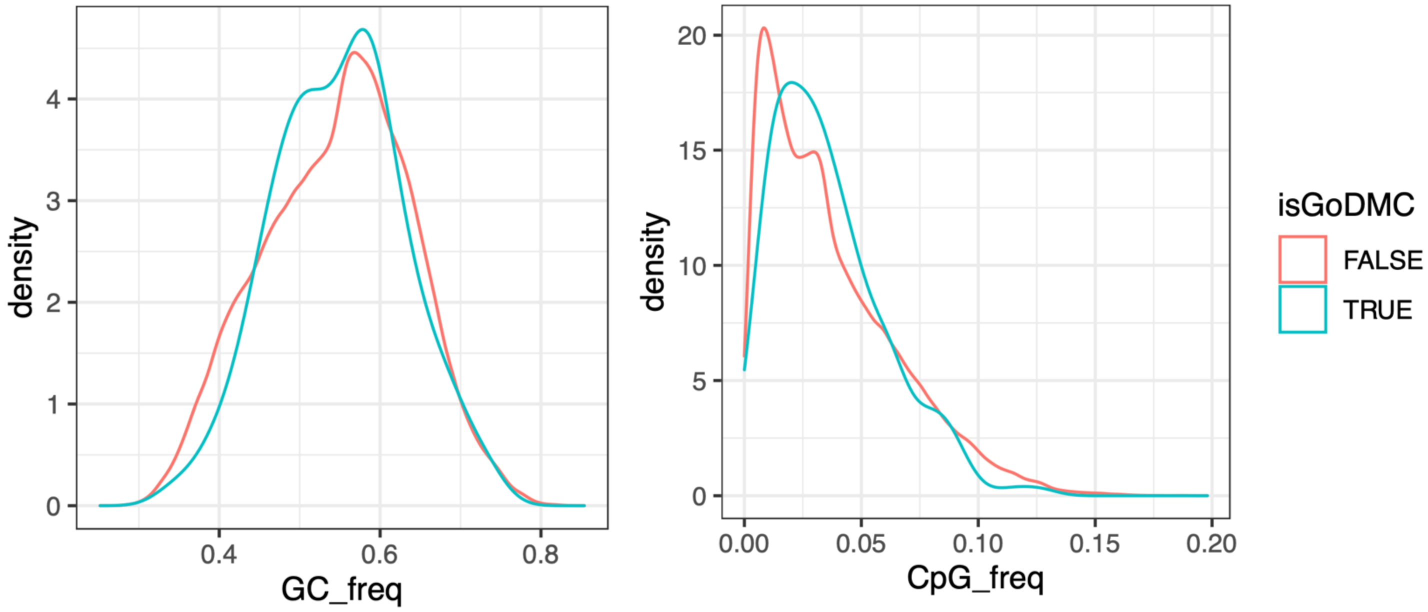


**Figure S6 |** Example of GC/CpG matching between DNAm sites of interest and background HumanMethylation450 DNAm sites for enrichment analysis in LOLA[2]. Is GoDMC true shown in blue (DNAm probes of interest), is GoDMC false shown in red (background HumanMethylatoin450 DNAm sites)

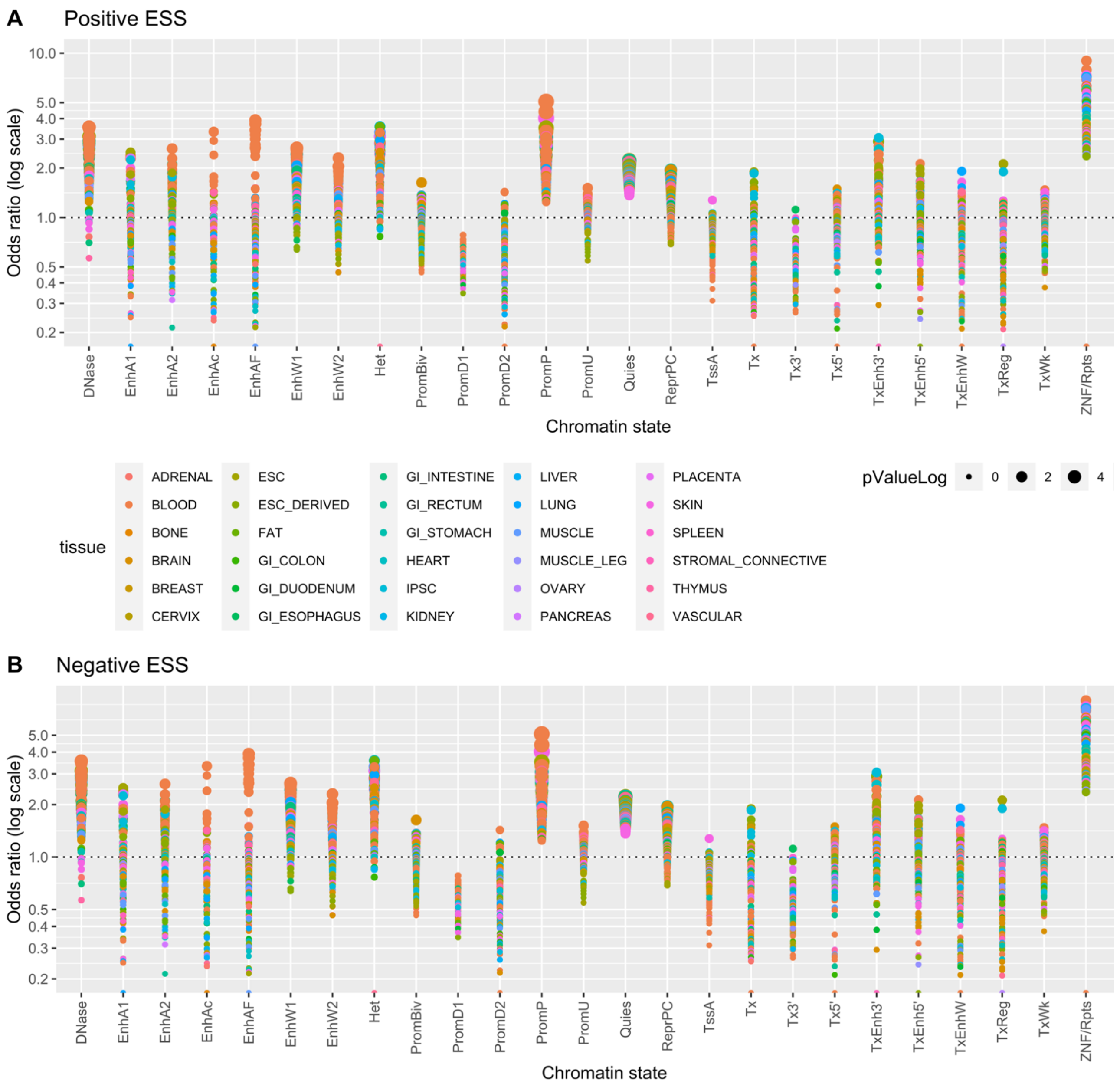


**Figure S7 | Enrichment or depletion of DNAm sites in predicted chromatin states for DNAm sites from the ESS probe set with estimates of *S* > 0.5 (positive ESS) and *S* < -0.5 (negative ESS).** Odds ratio (on log scale) shown on the *y* axis and chromatin state on the *x* axis. Size of circle represents the -log10 Pvalue. Enrichment analysis performed via one-sided Fisher's exact test implemented in LOLA[2]. 25 chromatin states abbreviations: TssA, Active TSS; PromU, Promoter Upstream TSS; PromD1, Promoter Downstream TSS with DNase; PromD2, Promoter Downstream TSS; Tx5', Transcription 5'; Tx, Transcription; Tx3', Transcription 3'; TxWk, Weak transcription; TxReg, Transcription Regulatory; TxEnh5', Transcription 5' Enhancer; TxEnh3', Transcription 3' Enhancer; TxEnhW, Transcription Weak Enhancer; EnhA1, Active Enhancer 1; EnhA2, Active Enhancer 2; EnhAF, Active Enhancer Flank; EnhW1, Weak Enhancer 1; EnhW2, Weak Enhancer 2; EnhAc, Enhancer Acetylation Only; DNase, DNase only; ZNF/Rpts, ZNF genes & repeats; Het, Heterochromatin; PromP, Poised Promoter; PromBiv, Bivalent Promoter; ReprPC, Repressed PolyComb, Quies, Quiescent/Low.

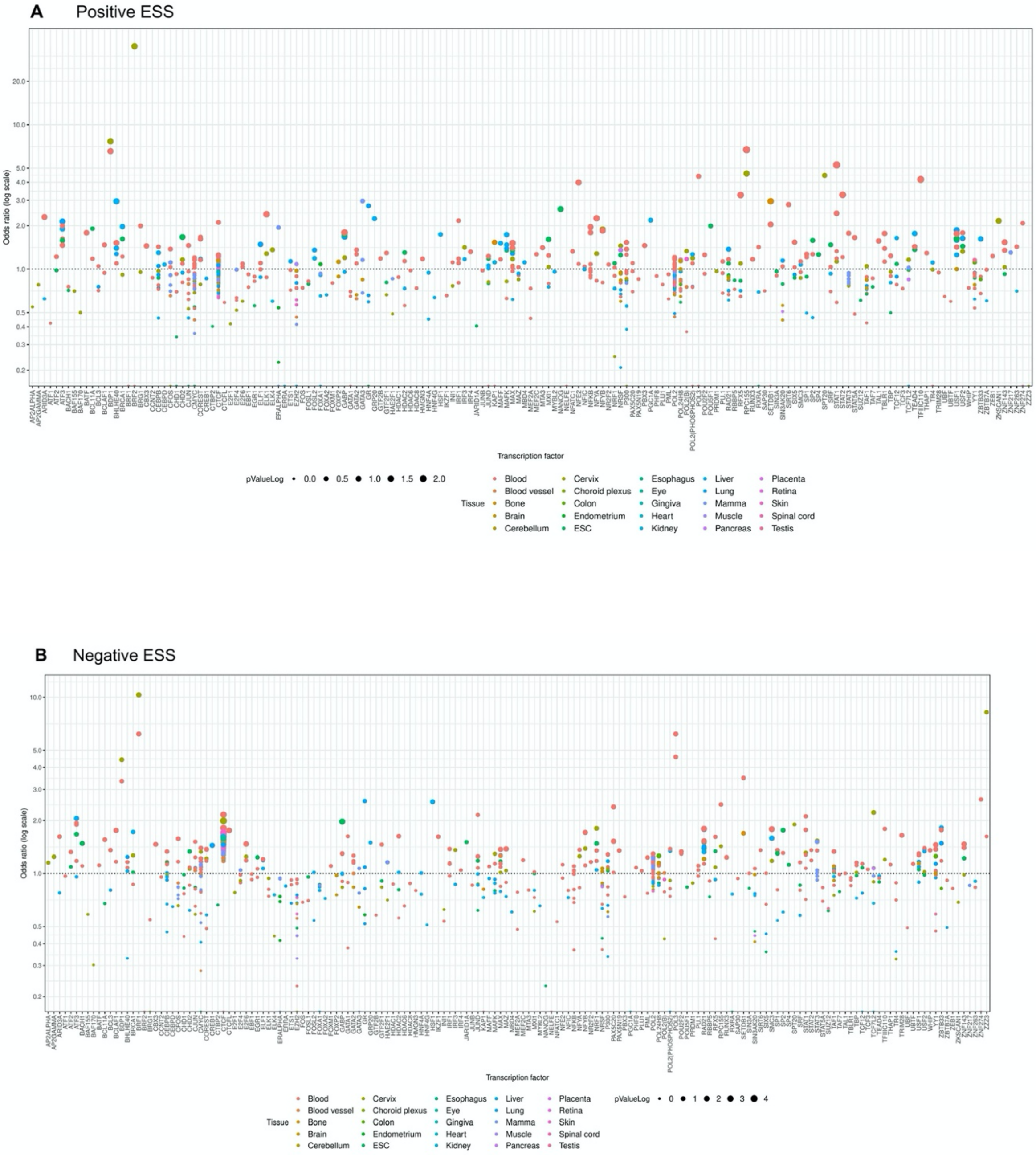


**Figure S8 | Enrichment or depletion of DNAm sites in transcription factors (TFs) for DNAm sites from the highly heritable probe set with estimates of *S* > 0.5 (positive highly heritable) and *S* < -0.5 (negative highly heritable).** Odds ratio (on log scale) shown on the *y* axis and chromatin state on the *x* axis. Size of circle represents the -log10 Pvalue. Enrichment analysis performed via one-sided Fisher's exact test implemented in LOLA[2].

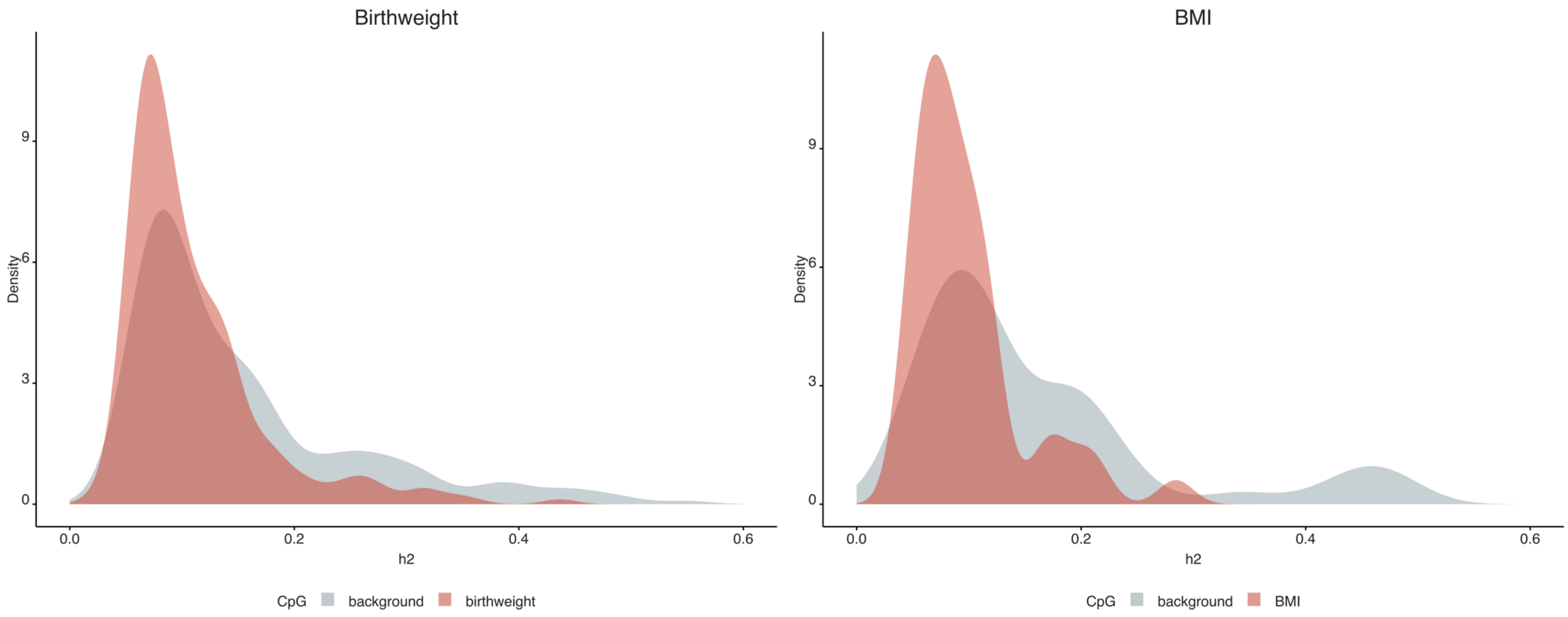


**Figure S9 | Distribution of estimates of $h_2$ for DNAm sites associated with birthweight and BMI compared to background DNAm sites.** Birthweight and BMI associated DNAm sites shown in red and matched DNAm sites (matched on GC/CpG content and heritability) shown in grey.

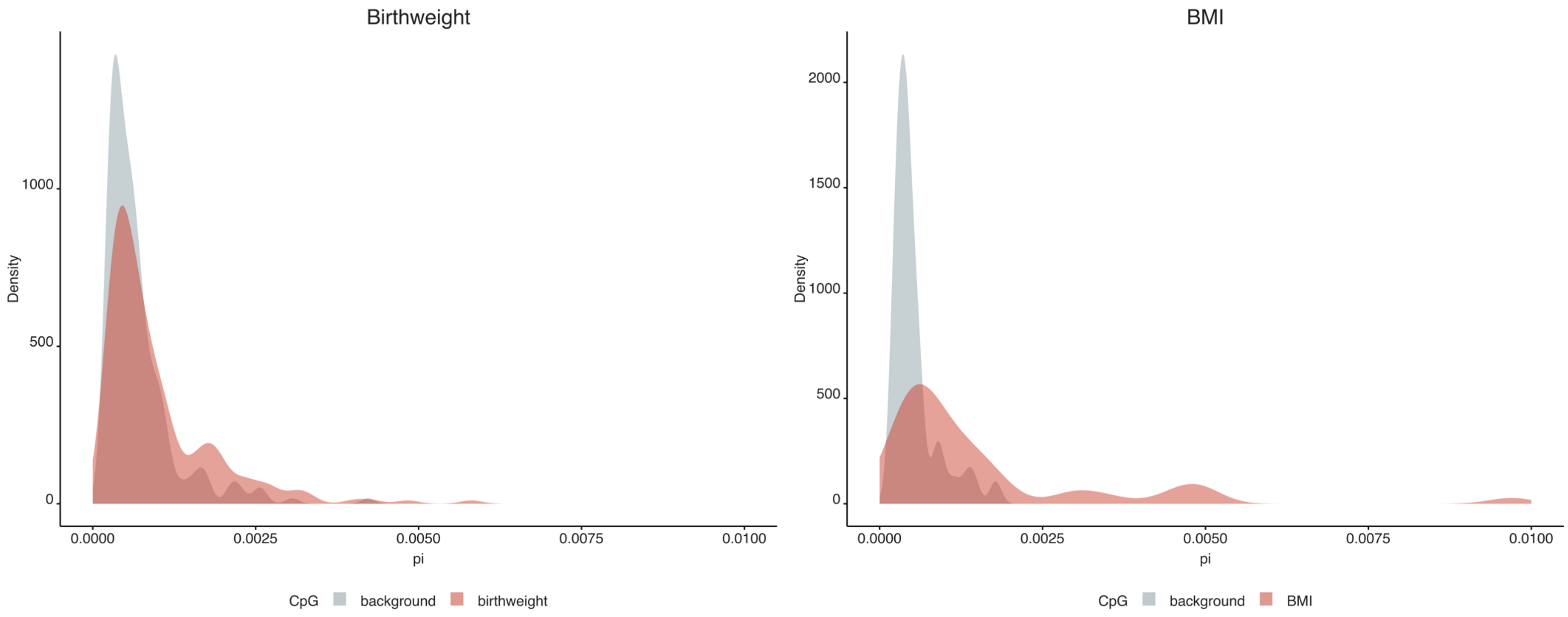


**Figure S10 | Distribution of estimates of $\pi$ for DNAm sites associated with birthweight and BMI compared to background DNAm sites**. Birthweight and BMI associated DNAm sites shown in red and matched DNAm sites (matched on GC/CpG content and heritability) shown in grey.
